# Generalizable and scalable protein stability prediction with rewired protein generative models

**DOI:** 10.1101/2025.02.13.638154

**Authors:** Ziang Li, Yunan Luo

**Affiliations:** School of Computational Science and Engineering, Georgia Institute of Technology

## Abstract

Predicting changes in protein thermostability caused by amino acid substitutions is essential for understanding human diseases and engineering proteins for practical applications. While recent protein generative models demonstrate impressive zero-shot performance in predicting various protein properties without task-specific training, their strong unsupervised prediction ability remains underexploited to improve protein stability prediction. We present SPURS, a deep learning framework that rewires and integrates two complementary protein generative models–a protein language model and an inverse folding model–and reprograms this unified framework for stability prediction through supervised fine-tuning on mega-scale thermostability data. SPURS delivers accurate, efficient, and scalable stability predictions with exceptional generalization to unseen proteins and mutations. Beyond stability prediction, SPURS enables broad applications in protein informatics, including zero-shot identification of functional residues, improved low-*N* protein fitness prediction, and systematic dissection of stability-pathogenicity for human diseases. Together, these capabilities establish SPURS as a versatile tool for advancing protein stability prediction and protein engineering at scale.

## Introduction

Thermodynamic stability, often quantified by changes in Gibbs free energy (Δ*G*), is a fundamental property of proteins. Understanding protein stability is crucial for engineering robust proteins for industrial and therapeutic applications ^1–3^. Although experimental techniques such as directed evolution have been successful in identifying stabilizing mutations, they require extensive experimental effort to screen numerous mutants, as stabilizing mutations are rare and the mutation exploration is often unguided. These constraints have driven the development of computational methods, particularly biophysical models and machine learning (ML) approaches ^4^, to predict stability changes (ΔΔ*G*) resulting from amino acid substitutions, offering cost-effective alternatives for engineering stabilized proteins.

While deep learning has revolutionized protein structure prediction through models like AlphaFold ^5^, no comparable transformative methods have emerged for protein stability prediction yet. This disparity stems primarily from data scarcity and the limitations of current computational models. Most current ML methods for stability prediction ^6–14^ are trained on relatively modest datasets ^8,9,15–18^, typically containing only hundreds to thousands of mutants across tens to hundreds of proteins. The mismatch between the limited data and the large demands of training data by modern ML models results in poor prediction generalization, particularly to unseen proteins or rare mutations. Although database consolidation efforts ^16,19,20^ have expanded data coverage, their inherent biases toward destabilizing variants, specific protein families, and experimental conditions continue to hinder progress in stability prediction, even as ML model complexity continues to grow.

Recent advances in mutagenesis experiments, such as complementary DNA (cDNA) proteolysis assays, offer new opportunities for ML-based protein stability prediction. A notable example is the mega-scale dataset (hereafter “Megascale” dataset) by Tsuboyama et al.^21^, which provides over 770,000 ΔΔ*G* measurements spanning all single and selected double amino acid variants across 479 diverse protein domains. This dataset, derived consistently from the same assay, represents an unprecedented resource for developing high-capacity ML models for protein stability prediction. However, fully realizing its potential requires ML models capable of capturing the underlying thermodynamic constraints driving protein folding and stability.

In parallel, protein generative models, including protein language models (pLMs) ^22–25^ and inverse-folding models (IFMs) ^26,27^, have recently emerged as “foundation models” for various ML tasks in protein informatics. Pre-trained to predict masked residue amino acids using context from unmasked residues (Fig. 1a), these models capture evolutionary patterns and biophysical constraints from vast natural protein sequence or 3D structure data. Studies have shown that these models can predict mutation effects in a zero-shot manner by calculating the log-likelihood ratios between mutants and wild-type proteins ^28–30^, correlating well with diverse protein fitness measures, including pathogenicity ^31^, binding affinity ^32^, and thermostability ^30,33^, even without task-specific supervised training.

**Figure 1:**
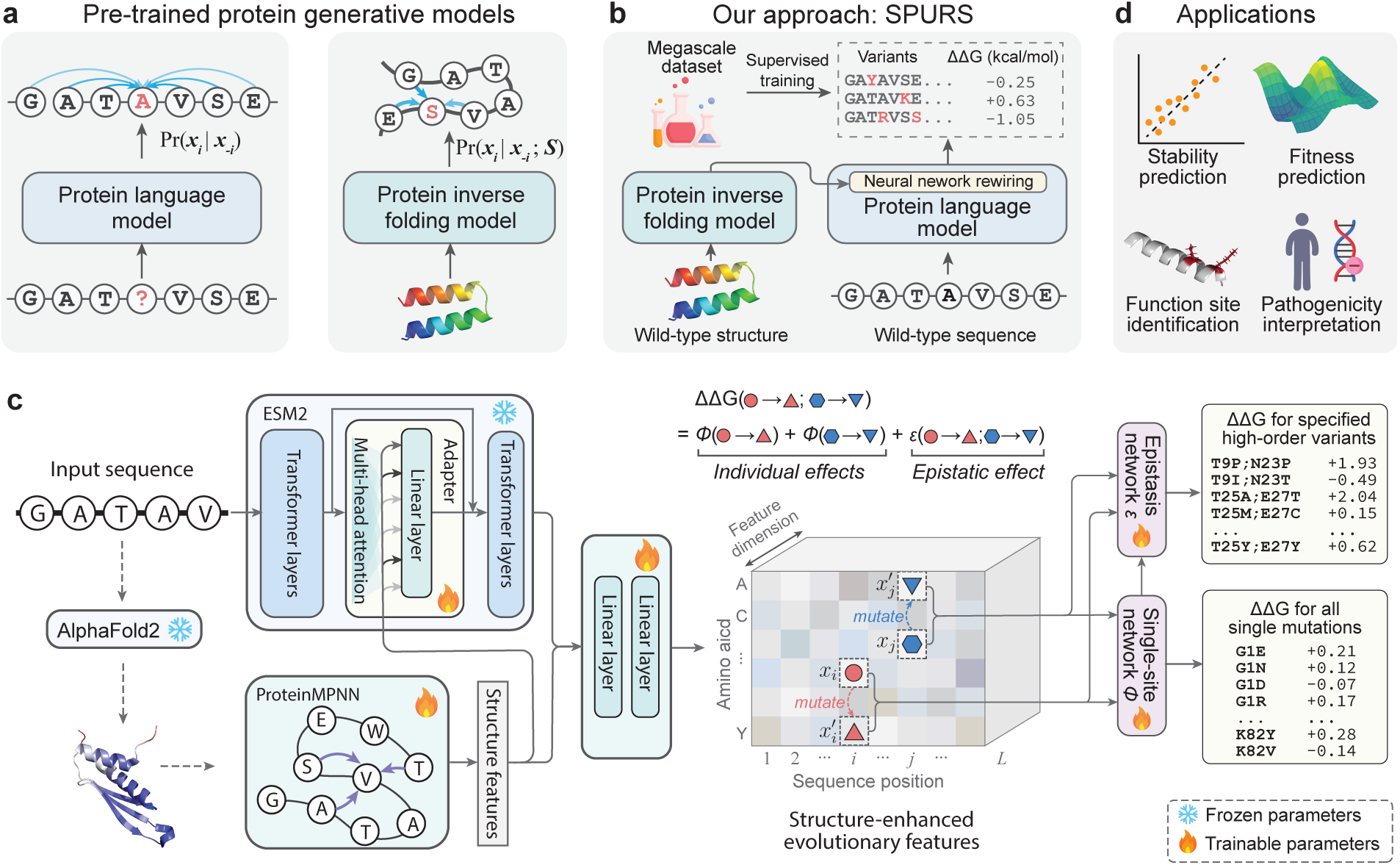
SPURS architecture. **a-b**, SPURS is a deep learning framework that rewires two pre-trained protein generative models (**a**) –a protein language model (ESM2) and an inverse folding model (ProteinMPNN)–to predict changes in protein thermostability (ΔΔ*G*) upon sequence mutations (**b**). **c**, The model takes as input a protein’s wild-type sequence and structure (predicted by AlphaFold2 if the experimental structure is unavailable). Through neural network rewiring, SPURS integrates evolutionary priors from ESM2 and structural features from ProteinMPNN to learn structure-enhanced evolutionary features. These representations are passed to a prediction module that efficiently decodes ΔΔ*G* predictions for all single mutations. The same features are reused by another decoder to predict epistatic effects of multiple mutations, enabling ΔΔ*G* prediction for higher-order mutants. **d**, SPURS’s performance is demonstrated in stability prediction and diverse applications in protein biology, including functional site identification, protein fitness prediction, and pathogenicity analysis. Abbreviations: Struct=Structure; Evo=Evolution.

These findings raise the natural expectation that combining pre-trained generative models with large-scale stability data through supervised training could yield improved stability prediction performance. However, recent studies reported that fine-tuning pLMs on the Megascale dataset often results in limited or even degraded performance compared to conventional ML models trained from scratch ^34,35^. One explanation of this under-performance lies in the sheer size of pLMs with ∼10^9^ parameters, making them prone to overfitting ^36^ despite the availability of mega-scale training data. Moreover, pLMs operate solely on linear amino acid sequences, lacking structural context of 3D residue interactions that are crucial for modeling protein thermostability.

Addressing this limitation, recent efforts have explored structure-aware generative models like Protein-MPNN ^26^ and ESM-IF ^37^ for stability prediction via fine-tuning ^38,39^. These models incorporated 3D structural information and have demonstrated improved generalization to unseen proteins, with some now recognized as state-of-the-art among ML-based stability predictors ^39,40^. However, IFMs like ProteinMPNN are intrinsically constrained by the quantity and diversity in their training data, which includes only experimentally determined structures (∼20k curated structures from the CATH database)–orders of magnitude fewer than the over 400M protein sequences in UniRef ^41^. While other IFMs like ESM-IF incorporate AlphaFold2-predicted structures to expand training coverage, this approach remains constrained by the number of structures that can be predicted and the accuracy of the predictions, failing to capture full sequence diversity. Consequently, IFMs underutilize rich evolutionary information from large-scale sequence data, while pLMs lack explicit structural grounding. We hypothesize that integrating pre-trained pLMs and IFMs—trained on complementary modalities—could synergistically improve stability prediction by combining sequence-derived evolutionary priors with structure-based geometric features. However, fusing these models presents substantial challenges due to differences in their data modality, architectural design, and training scale.

In this work, we introduce SPURS (stability prediction using a rewired strategy), a deep learning framework that integrates sequence- and structure-based protein generative models for generalizable stability prediction (Fig. 1b). At the core of SPURS is a model rewiring strategy that implants a lightweight Adapter module ^42,43^ into a pLM (ESM ^24^), enabling it to incorporate structure-informed features learned by an IFM (ProteinMPNN ^26^) while preserving the pLM’s evolutionary priors learned from large-scale sequence training (Fig. 1c). This design captures both IFM’s features specific to structural stability alongside pLM’s features supporting broad generalization. It also allows parameter-efficient fine-tuning, requiring updates to only a small fraction of trainable parameters to jointly model sequence and structure for ΔΔ*G* prediction. Importantly, SPURS is highly scalable by design: it predicts ΔΔ*G* values for all possible point mutations of a protein in a single forward pass of the neural network, in contrast to many prior methods that require separate evaluations per mutant. This scalability enabled us to perform large-scale stability predictions across the human proteome.

SPURS demonstrates strong generalization in protein stability prediction. We trained SPURS on the Megas-cale dataset and systematically benchmarked performance across 12 diverse datasets measuring thermostability (ΔΔ*G*) and melting temperature changes (Δ*T_m_*). SPURS consistently outperformed state-of-the-art methods, achieving robust generalization to unseen mutations and proteins across datasets. In particular, SPURS excels at identifying stabilizing mutations, a persistent challenge for most existing methods due to the pronounced imbalance toward destabilizing variants in current stability datasets ^44^.

Beyond stability prediction, SPURS enables a range of compelling applications in protein biology (Fig. 1d). We show that, when paired with a pLM, SPURS can accurately identify functionally important residues involved in protein binding using only sequence input, without relying on binding structure information. We also demonstrate that SPURS’s ΔΔ*G* predictions serve as informative priors for guiding the prediction of protein variant fitness, enabling a simple yet effective regression model that enhances low-*N* fitness prediction. Finally, leveraging its scalability, we applied SPURS to the human proteome to dissect the role of stability loss in variant pathogenicity and uncover the distinct molecular mechanisms of different pathogenic mutations. Together, these capabilities position SPURS as a versatile method for protein stability modeling, with broad applications in structural biology, protein engineering, and functional genomics.

## Results

### SPURS: Thermostability prediction leveraging protein generative models

SPURS is a deep learning framework designed to predict changes in protein thermostability (ΔΔ*G*) resulting from amino acid substitutions (Fig. 1). Given a wild-type protein sequence, SPURS predicts ΔΔ*G* for all possible point mutations or specified higher-order mutations. To inform these predictions, SPURS explicitly incorporates the 3D structure of the wild-type protein as additional input. Structure information is represented as atomic coordinates and abstracted into a graph with atoms as nodes and edges defined by spatial proximity (Methods). When an experimental structure is unavailable, SPURS employs AlphaFold ^5^ to predict the structure, ensuring broad applicability.

At the core of SPURS is an effective integration of two pre-trained protein generative models (Fig. 1a): ESM2^24^, a Transformer-based protein language model (pLM) trained on protein sequences, and Protein-MPNN ^26^, a graph neural network-based inverse-folding model (IFM) trained on protein structures. SPURS utilizes ProteinMPNN as a structure encoder to extract geometric features important for protein stability while leveraging sequence evolutionary priors learned by ESM to dissect mutation effects on stability (Methods).

Rather than treating these two models separately or integrating their outputs heuristically, SPURS employs Adapter ^42,43^, a lightweight neural network module, to rewire ProteinMPNN’s structure-derived embeddings into ESM’s sequence embeddings, learning structure-enhanced sequence representations in a parameter-efficient fashion (Fig. 1b-c; Methods). During training, only Adapter and ProteinMPNN parameters are updated, while ESM’s parameters remain fixed. This integration strategy introduces only minimal architecture alterations to ESM and ProteinMPNN, preserving their rich evolutionary priors learned from pre-training while avoiding overfitting. This approach also makes SPURS highly data-efficient for fine-tuning, as it dramatically reduces trainable parameters by 98.5% compared to full ESM fine-tuning used in previous studies ^35,39^. Although this work specifically uses ESM and ProteinMPNN, SPURS is a model-agnostic framework and can integrate other sequence- and structure-based generative models ^25,37^ for stability prediction.

SPURS offers significant scalability advantages through its algorithmic design. At the output layer, SPURS predicts ΔΔ*G* for all possible point mutations for the input protein in a single forward pass of the neural network (Fig. 1c). This represents a major advancement over existing stability prediction methods ^10,34,35,45,46^, which typically require separate forward passes for each mutant sequence, resulting in computational cost of O(*L* × 20) forward passes for a protein of length *L*. In contrast, SPURS transforms this one-mutant-per-pass approach to an all-mutants-per-pass paradigm. Rather than processing mutant sequences as input, SPURS conditions on wild-type sequence and structure to predict stability changes for all possible point substitutions simultaneously. This is achieved by learning per-residue latent representations and using a decoder shared across all residues to predict effects of substituting each residue with any of the 20 amino acids (Methods). This all-at-once inference paradigm reduces the required number of forward passes from O(*L* × 20) to O(1), enabling efficient large-scale protein stability profiling. In our experiments, SPURS predicted ΔΔ*G* for all single mutants across 118 full-length proteins from the ProteinGym benchmark ^29^ (mean length: 492 residues) in under 20 seconds on a single NVIDIA A40 GPU.

SPURS was trained on the Megascale dataset, comprising over 200,000 ΔΔ*G* measurements for single amino acid substitutions across over 200 proteins. This unprecedented data size enables SPURS to learn generalizable representations for unseen proteins and mutations. Additionally, Megascale’s dense mutational coverage–with each wild-type protein exhaustively mutagenized for all possible point mutations–enables SPURS to learn fine-grained, generalizable patterns across protein space and effectively capture the full ΔΔ*G* landscape of *L* × 20 possible substitutions.

We further extended SPURS to predict stability for higher-order mutants by modeling ΔΔ*G* of a multi-mutation variant as the sum of individual mutation effects plus an epistatic term capturing non-additive interactions. SPURS reuses point-mutation predictions, inferred from a single forward pass, and employs an additional lightweight decoder network to predict epistatic effects (Methods). This extended SPURS framework was trained on all single- and double-mutation ΔΔ*G* measurements from Megascale. This design captures complex combinatorial effects while maintaining efficiency. Together, SPURS provides a scalable, generalizable framework for stability prediction across both single and combinatorial mutation regimes.

### SPURS enables accurate and generalizable protein stability prediction

To evaluate SPURS’s performance in predicting protein stability changes upon mutations, we curated 12 datasets of stability measurements from published studies (Methods). These datasets vary in protein diversity and mutant coverage (Fig. 2a), collectively forming a comprehensive benchmark for assessing model accuracy and generalization. We first focused on the Megascale dataset ^21^, using the training, validation, and test splits (8:1:1 ratio) established in the ThermoMPNN study ^38^, comprising ΔΔ*G* measurements for 272,721 single-substitution mutants across 298 proteins. To prevent information leakage, we removed from the training and validation splits any sequences with *>* 25% sequence identity to those in the test split or other independent datasets (Methods). Throughout our experiments, SPURS was trained on this filtered Megascale training set.

**Figure 2:**
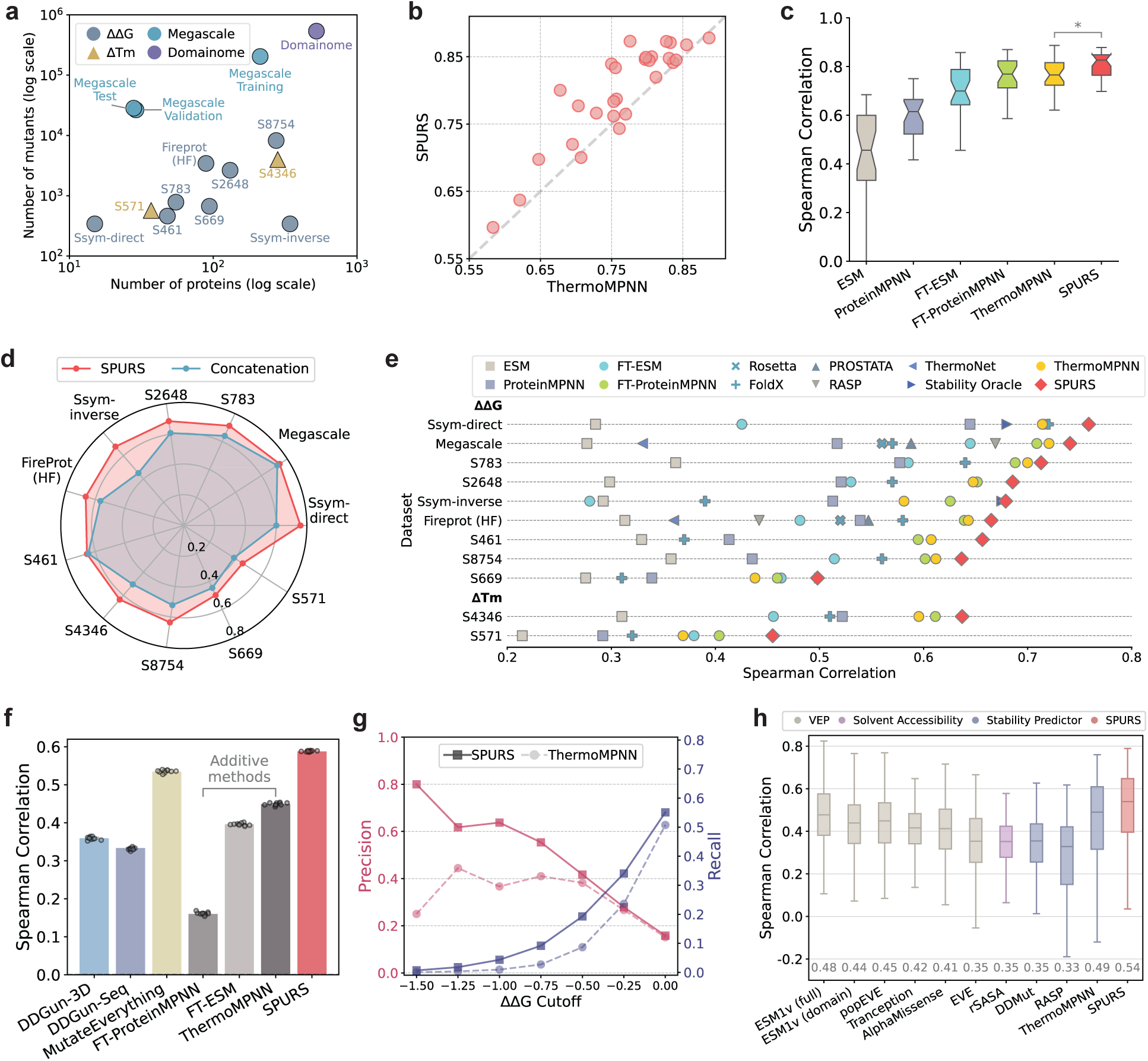
SPURS achieves accurate prediction for protein stability. a, The numbers of proteins and mutants across the datasets used for training and evaluation. **b,** Performance comparison of SPURS and ThermoMPNN (a state-of-the-art deep learning-based stability model) on the Megascale test set. Each dot represents the Spearman correlation of SPURS and ThermoMPNN on a single protein test set. **c,** Performance comparison between SPURS and additional baseline methods on the Megascale test set (*n* = 28). ESM and ProteinMPNN are evaluated in a zero-shot setting using sequence likelihood differences between mutant and wild-type; FT-ESM and FT-ProteinMPNN denote fine-tuned (FT) versions of ESM and ProteinMPNN trained on the same Megascale training split as SPURS and ThermoMPNN. *: two-side Mann-Whitney U test *P* =0.003. **d,** Effectiveness of SPURS’s rewiring strategy for integrating sequence- and structure-based protein generative models. SPURS was compared to a straightforward integration approach that directly concatenates the final-layer embeddings of ProteinMPNN and ESM (“Concatenation”). **e,** Performance of SPURS versus baseline methods across the Megascale test set and ten independent benchmarks. **f,** Performance for predicting ΔΔ*G* of double-mutation mutants in the Megascale test set. Bar plots represent mean*±*s.d. of the performances on ten 80% bootstrapped subsets (*n* = 10) from all double mutants in the Megascale test split. DDGun-3D and DDGun-seq use structure and sequence features to predict ΔΔ*G*, respectively. FT-ESM, FT-ProteinMPNN, and ThermoMPNN are single-mutant predictors extended to predict for double mutants by simply adding the predicted ΔΔ*G* of constituent individual mutations (“additive methods”); other methods directly support combinatorial predictions. **g,** Precision–recall comparison between SPURS and ThermoMPNN for identifying stabilizing mutations in the Megascale test set. **h,** Performance comparison on the Domainome dataset (*n* = 522). ESM1v (full) takes the full-length protein sequence as input, whereas ESM1v (domain) uses domain subsequences. Abbreviations: VEP = Variant effect predictors; rSASA = Relative solvent-accessible surface area. Box plots in **c** and **h** indicate the median (center line), 25th–75th percentiles (box), and whiskers represent data within 1.5*×* the interquartile range from the box.

We first compared SPURS to ThermoMPNN, the state-of-the-art ML model for stability prediction, on the Megascale test set covering 28,312 mutants across 28 proteins. SPURS outperformed ThermoMPNN in Spearman correlation (median 0.83 vs. 0.77, Mann-Whitney U test *P <* 0.05) in 24 out of 28 proteins (Fig. 2b) and remained comparable for the remaining four proteins. This improvement can be attributed to SPURS’s effective integration of both sequence and structural priors from ESM and ProteinMPNN, in contrast to ThermoMPNN, which fine-tuned structure-based ProteinMPNN. Further, SPURS’s rewiring strategy outperformed a straightforward integration approach that simply concatenates final-layer embeddings from ESM and ProteinMPNN across all test datasets (Fig. 2d), suggesting the consistent effectiveness of its integration.

Another ablation study, where we fine-tuned ESM and ProteinMPNN individually using multi-layer perceptrons (MLPs) on top of frozen ESM or ProteinMPNN layers, confirmed that fine-tuning either model alone led to performance drops compared to their integration in SPURS (Fig. 2c). Interestingly, even sequence likelihoods predicted by unsupervised ESM and ProteinMPNN exhibited a non-trivial Spearman correlation with ΔΔ*G* measurements in Megascale (0.46 and 0.61, respectively). This observation is consistent with previous studies ^30^, suggesting that protein generative models, even without explicit training on stability data, capture evolutionary features predictive of stability. This forms the basis of our hypothesis and other works ^35,38^ that supervised fine-tuning of protein generative models improves stability prediction. Next, we evaluated SPURS’s generalizability using eight independent test sets from other studies ^8,9,15,18,38,45,47^.

We excluded sequences in the Megascale training and validation sets that had more than 25% sequence identity with these test sets. We additionally included six leading baselines (Supplementary Note B.1), including biophysical models (FoldX ^48^, Rosetta ^49^) and ML methods (PROSTATA ^10^, RASP ^34^, Stability Oracle ^50^, ThermoNet ^7^, ThermoMPNN ^38^). SPURS showed significantly higher (for 7/8 test sets) or comparable Spearman and Pearson correlations across all datasets compared to these baselines (Fig. 2e and Supplementary Tables 1, 2, and 3).

We also explored SPURS’s ability to generalize to melting temperature (Δ*T_m_*) prediction, another measure of protein stability. Even though SPURS was only trained on ΔΔ*G* data, it demonstrated improved Spearman correlations on two Δ*T_m_* datasets ^45^, S4346 and S571 (Fig. 2e), which highlights SPURS’s broad capability to capture stability-related features beyond its training data.

To assess SPURS’s ability to predict ΔΔ*G* for higher-order mutants, we evaluated its performance on double-mutation data from the Megascale test set. SPURS outperformed DDGun ^14^ and MutateEverything ^51^, two competitive existing methods that support stability prediction for combinatorial mutations (Fig. 2f), even though advantages were given to MutateEverything as 13 of 20 proteins in our test set were included in its training set. In the comparison, we also included methods that only predict stability change for point mutations, including the fine-tuned ESM, ProteinMPNN, and ThermoMPNN, to predict for high-order mutants through additive effects of constituent individual mutations. Results showed that SPURS substantially out-performed these additive approaches (Fig. 2f), suggesting that multiple mutation effects on stability are not simply additive and that SPURS’s modeling of epistatic effects is critical to achieving the performance gain.

Stabilizing mutations are of particular interest for protein engineering but are often rare and underrepresented in most datasets. As a result, many models tend to optimize prediction accuracy for overrepresented destabilizing variants, inflating apparent accuracy ^50,52^. To assess robustness in this regime, we evaluated SPURS’s ability to prioritize stabilizing mutations (defined as ΔΔ*G<* − 0.5 kcal/mol ^53^; *N* =1,178) from a much larger pool of destabilizing ones (*N* =27,139) in the Megascale test set. Across various prediction thresholds, SPURS consistently outperformed ThermoMPNN in both precision and recall (Fig. 2g), indicating its effectiveness in identifying stabilizing mutations.

Finally, we applied SPURS to the recently released Human Domainome dataset ^40^, which quantifies the impact of human missense variants on protein stability using protein abundance in cells. This dataset contains 563,534 variants across 522 proteins, offering the largest diversity and coverage among our benchmarks (Fig. 2a). The original Domainome study reported that ThermoMPNN outperformed several other models for stability prediction, including general variant effect predictors (e.g., AlphaMissense ^54^ and EVE ^31^), structural features (relative solvent accessibility), and dedicated stability predictors. We thus re-examined this evaluation and additionally compared SPURS with these reported methods. The result showed that SPURS significantly improved upon the best baseline, ThermoMPNN (correlation 0.54 vs. 0.49), further demonstrating SPURS’s generalizability across large-scale variant effect landscapes of stability (Fig. 2h).

Taken together, our benchmark results demonstrate that SPURS achieved state-of-the-art performance for protein stability prediction, with superior generalizability and reduced bias compared to existing models.

### SPURS identifies functionally important sites in proteins

Proteins perform diverse cellular functions, largely through interactions with other molecules. Identifying the residues responsible for these interactions is key to understanding molecular mechanisms and developing biomedical applications. Stability is just one biophysical property that contributes to protein function, while others, such as binding specificity and enzymatic activity, often trade off with stability during evolution ^55^. Thus, the loss of function due to mutations can be attributed to either direct disruption of molecular interactions or structure destabilization that leads to reduced protein abundance. Mutations at protein binding interfaces, active sites, and allosteric sites frequently have larger effects on function than what stability changes alone can explain ^21,56,57^, making it complicated to deconvolve the effects of substitutions on intrinsic function from those on stability ^57,58^. Recent experimental studies attempted to resolve this biophysical ambiguity by quantifying mutation effects on both protein binding and abundance, allowing comprehensive mapping of functional sites ^56,59^. Inspired by this, we hypothesized that a similar strategy, combining SPURS’s stability predictions with evolutionary fitness scores from pLMs, could help disentangle mutation effects on function and identify functional sites.

Specifically, we used SPURS to predict ΔΔ*G* and ESM1v ^28^ to estimate the evolutionary fitness of a protein variant (Methods). Here, ‘fitness’ broadly refers to protein functions like binding affinity, catalytic activity, and more. ESM1v has been shown to be effective for zero-shot predictions of mutation effects on protein fitness ^28^. We fit a sigmoid function to model the non-linear relationship between stability and fitness (Fig. 3a; Methods), following prior work that employed non-linear Boltzmann distribution to model the relationship between free energy changes caused by mutations in protein folding and those in protein binding ^56,59–62^. A recent study showed that the residuals (fitting deviation) from the fitted sigmoid curve indicate whether mutations have larger or smaller effects on protein fitness than can be explained by changes in stability ^40^. Our approach generalized their study by extending functional site identification beyond the restricted set of 500 protein domains with experimental stability data ^40^, scaling up to diverse, full-length proteins using SPURS’s accurate stability predictions. We computed the fit residuals for all single mutations in a given protein (Fig. 3b) and defined a per-site function score by averaging residuals across all mutations at each site (Methods). Residues with high function scores are likely to be functionally important ^40^ (Fig. 3b,c).

**Figure 3:**
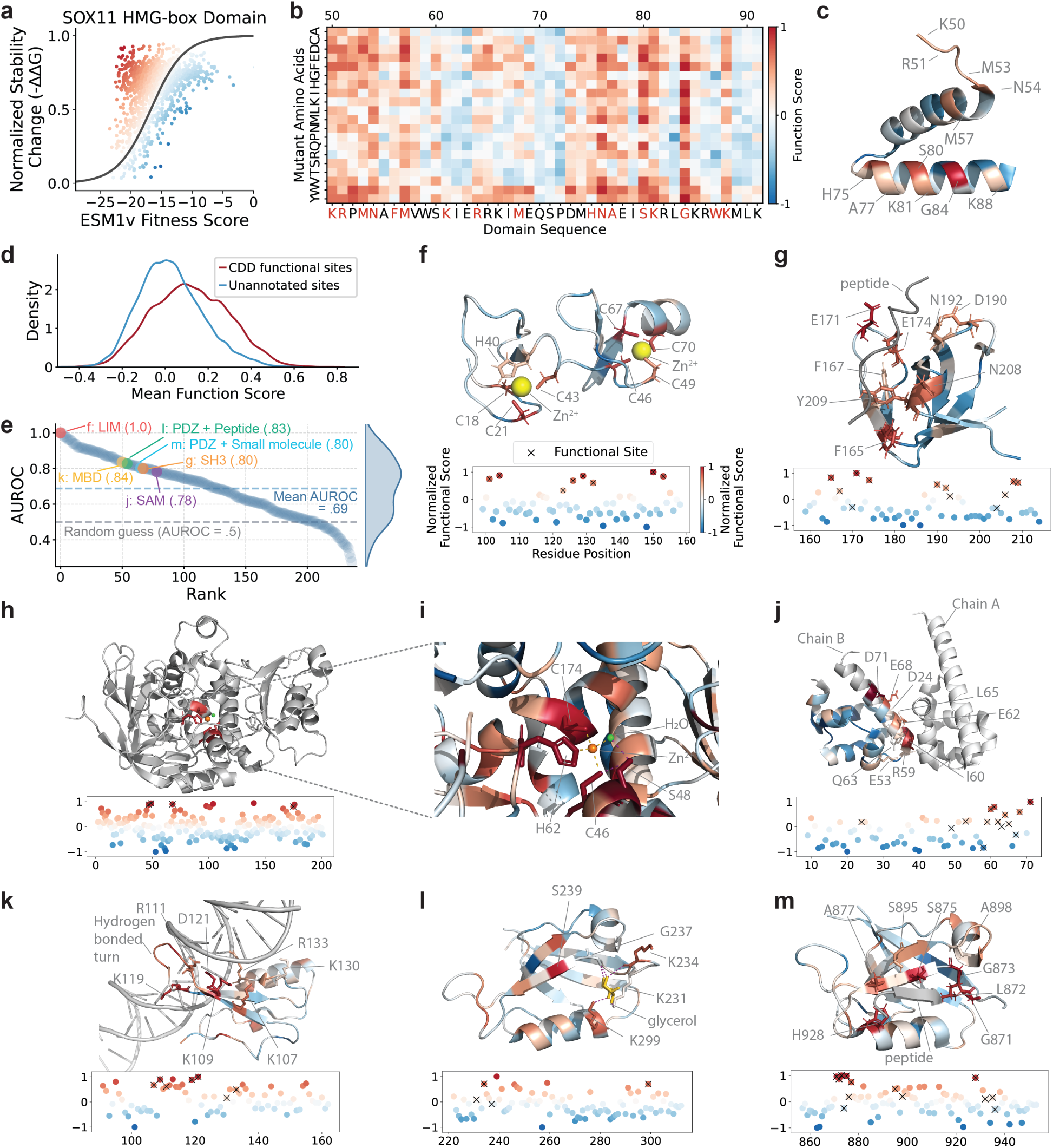
SPURS accurately guides functional site annotation. **a**, ESM1v-predicted fitness scores and min-max normalized stability changes (-ΔΔ*G*) predicted by SPURS for the HMG-box domain (UniProt ID: P35716). A sigmoid function is fitted to model the relationship between fitness and normalized stability change. Color gradient represents SPURS’s predicted function score. **b**, Heatmap depicting function scores of the HMG-box domain, with red letters indicating CDD-annotated DNA-binding sites. **c**, Structure of the HMG-box domain (PDB ID: 6T7C), where residues are colored based on SPURS’s predicted function scores and DNA-binding sites are annotated. **d**, Distribution of SPURS’s predicted function scores for functional sites annotated in CDD versus other sites across 239 human domains. **e**, SPURS’s AUROC performance for functional site prediction across 239 human domains. **f-m**: The structures colored by SPURS’s function scores for the six domains highlighted in **e**. Below each structure, a scatter plot shows the predicted function score for each residue, with CDD-annotated functional sites marked as *×*. **f**, LIM domain in FHL1 (UniProt ID: Q13642; PDB ID: 1X63). **g**, SH3 domain in GRB2 (P62993, 1IO6). **h**, Alcohol dehydrogenase (P00327, 1QLH). **i**, Zoom-in view of the zinc ion around the catalytic center. **j**, SAM domain in CNKSR2 (Q8WXI2, 3BS5). **k**, MBD domain in MECP2 (P51608, 5BT2), **l**, PDZ domain in DLG3 (Q92796, 2FE5), **m**, PDZ domain in SCRIB (Q14160, 6XA7).

We evaluated this approach for identifying function sites using 239 proteins from the Domainome dataset ^40^ with residue-level functional site annotations in the Conserved Domain Database (CDD) ^63^. Among 14,434 residues across these proteins, 3,516 were labeled as functional in CDD, while the rest were considered non-functional. SPURS’s function score significantly distinguished functional from non-functional sites (Fig. 3d; *t*-test *P <* 1 × 10^−1^^50^). At the individual protein level, SPURS achieved an average AUROC of 0.69 (Fig. 3e). Given that SPURS was not trained on functional site labels in contrast to previous supervised methods ^64^, these results demonstrated its strong unsupervised capability for identifying functional sites and scaling beyond with experimental stability data.

We further explored SPURS’s predicted functional sites across seven protein domains that ranked high or mid in AUROC (Fig. 3e). These proteins were chosen to cover human domains that represent various sizes (56–97 residues), diverse structural folds, and different functions, including protein-protein interaction, DNA binding, ion interaction, and enzymatic activity. By mapping SPURS’s function score onto their 3D structure, we found high-score regions significantly enriched for functional sites (Figs. 3f-m).

In the LIM domain, which contains two four-residue zinc fingers, SPURS assigned high scores exclusively to the zinc-coordinating residues (Fig.3f). Similarly, in the SH3 domain, SPURS accurately captured its critical peptide-binding sites (Figs. 3g). We extended this analysis to proteins not presented in Domainome, such as alcohol dehydrogenase (EC 1.1.1.1), a zinc-dependent enzyme. SPURS successfully identified its key zinc-binding residues (C46, H62, and C174) and a stabilizing residue (S48), which forms hydrogen bonds critical for the enzyme’s structure (Fig. 3h-i)

Notably, despite being trained only on single-chain inputs, SPURS identified interaction interfaces and binding sites. For example, it highlighted SAM domain residues near the heterodimer interaction interface (Fig. 3j), even though its interacting partner’s sequence or structure was not provided as input to SPURS. A similar observation was made for the MBD domain, where SPURS not only recovered DNA binding sites but also identified a *β*-turn located between two *β*-strands and close to the DNA helix (Fig. 3k). While this *β*-turn was not annotated by CDD as functional sites, it appeared to coordinate the protein-DNA binding, suggesting the potential of SPURS for discovering new functional sites.

SPURS also demonstrated consistency across different proteins harboring the same domain. For example, in two proteins with PDZ domains (Figs. 3l-m), SPURS assigned high function scores to several residues that occupy structurally corresponding positions in both proteins. A serine residue (S239 in Fig. 3l and S875 in Fig. 3m) consistently received the highest function score in both structures despite their different structural contexts. While SPURS prioritized consistent functional sites for both proteins, it also captured the context-dependent nature of binding and gave higher scores to residues involved in specific ligand interactions unique to each protein. In one case where the domain binds to a small glycerol molecule (Fig.3l), a lysine (K299) interacting with the molecule received a higher score than its counterpart in the other protein, while in the second case where the same domain binds to a peptide (Fig.3m), SPURS prioritized peptide-proximal residues (L872, H928). Importantly, the ligands were only shown for visualization and were not provided as input to SPURS. Nonetheless, SPURS was able to identify both conserved functional sites for the same domain across proteins and context-dependent sites specific to each protein’s binding function.

These case studies illustrate SPURS’s strong agreement with CDD annotations and its ability to identify both conserved and context-specific functional sites. Unlike previous methods that rely on docking structures ^65^ or supervised training ^64^, our approach is unsupervised, requiring only sequence input, with structural predictions generated by AlphaFold when needed. This approach alleviates data bottlenecks in functional site annotation, making it a powerful tool for applications like hotspot identification in protein engineering.

### SPURS improves low-*N* protein fitness prediction

Having established SPURS’s accuracy in predicting protein stability changes, we explored whether it could extend to enhancing the prediction of mutation effects on broader protein properties beyond stability. Many laboratory assays have been developed to measure various protein properties like binding affinity, expression, and solubility, often generally referred to as fitness. However, experimental techniques can only probe a tiny fraction of the exponentially large sequence space and screen their fitness, making it critical in protein engineering to develop ML models that generalize well from small-sized (low-*N*) fitness data to predict for unseen sequences ^27,36,66^.

Here, we aim to improve low-*N* fitness prediction models with SPURS. Proteins need to be structurally stable to perform functions. We thus hypothesized that SPURS’s stability predictions could serve as informative priors for fitness prediction. We propose a simple yet effective approach that incorporates SPURS’s ΔΔ*G* prediction to improve protein fitness prediction. Our approach was inspired by a leading supervised low-*N* fitness prediction model called ‘Augmented model’ ^27^. To predict the fitness of a protein variant, the Augmented model uses as input features the one-hot-encoded sequence and an evolutionary density score, which is the likelihood ratio between the mutant and the wild-type sequences predicted by pLMs like ESM or other sequence density models (e.g., DeepSequence ^67^ or EVE ^31^), to train a Ridge regressor on fitness data (Fig. 4a). We extended the Augmented model by incorporating SPURS’s ΔΔ*G* predictions as an additional feature in the regressor (Fig. 4a; Methods), yielding an enhanced model denoted as ‘SPURS-augmented ESM’ when ESM is used as the sequence density model, or similarly if other models are used.

**Figure 4:**
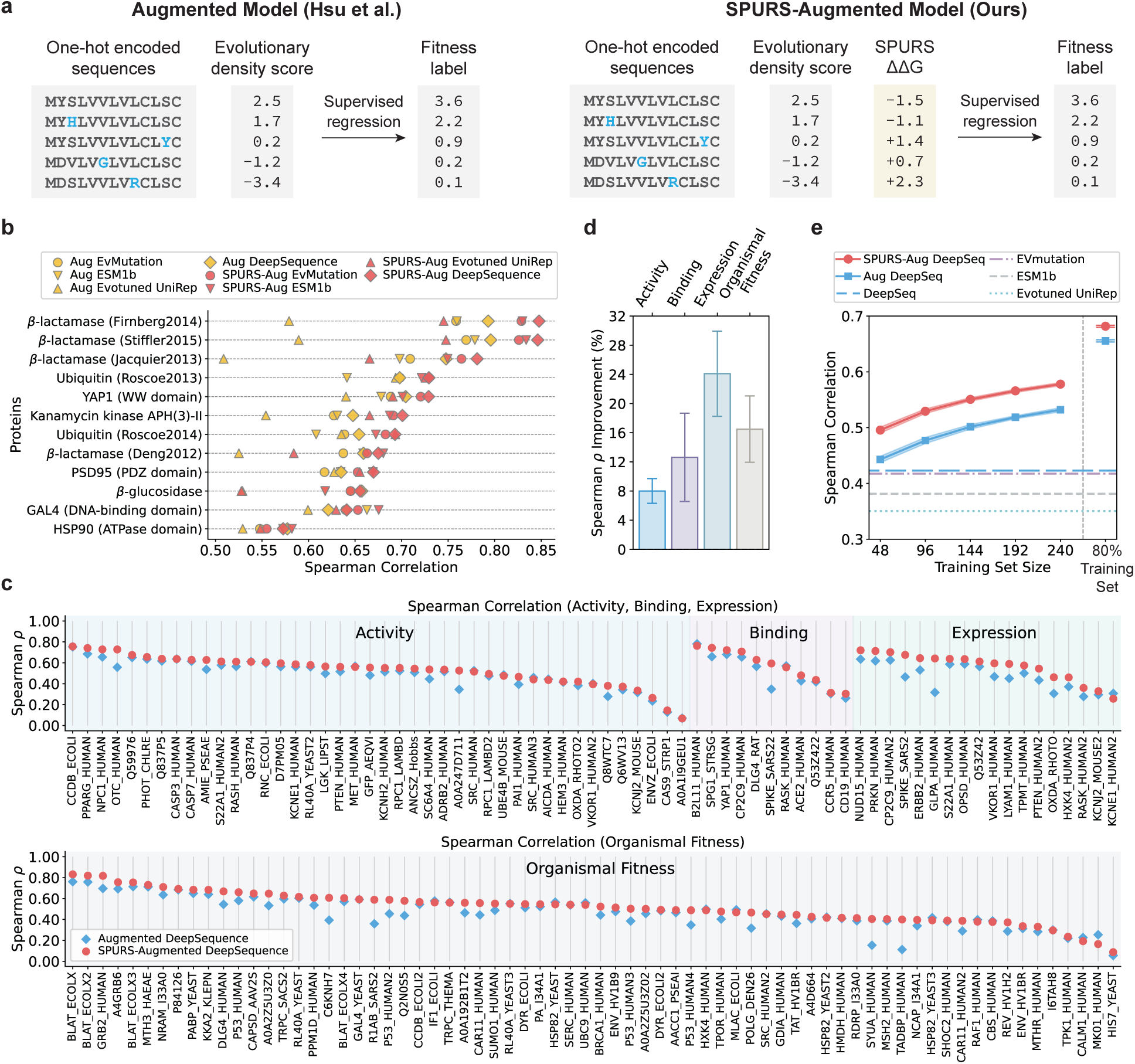
SPURS-augmented models enhances low-*N* protein fitness prediction. **a**, Architecture of the SPURS-augmented model. Compared to the original Augmented models ^27^, our approach adds SPURS’s predicted stability change as an additional feature. **b**, Performance comparison between SPURS-augmented models (squares) and their corresponding Augmented models (dots) on 12 proteins benchmarked in Hsu et al. ^27^. Aug=Augmented. **c**, Performances of SPURS-augmented DeepSequence and Augmented DeepSequence across 141 DMS datasets in ProteinGym, excluding stability-related sets and those with fewer than 96 variants. DMS dataset names on x-axis were abbreviated, with a mapping to their full names defined by ProteinGym provided in Supplementary Table 5. Results in **b-c** show the mean Spearman correlation over 20 repetitions of randomly sampled training sets. **d,** Relative Spearman correlation improvement of SPURS-augmented DeepSequence over Augmented DeepSequence, grouped by fitness categories from **c** (Activity, n = 42; Binding, n = 11; Expression, n = 18; OrganismalFitness, n = 70). **e,** Low-*N* prediction performance comparison between SPURS-augmented DeepSequence and Augmented DeepSequence across varying training set size *N*, from 48 variants up to 80% of each dataset. In **d-e**, plots show the mean*±*s.d. of the test performances of 20 repetitions of model training, with different random seeds used to randomly sample training data.

We compared SPURS-augmented models against a series of Augmented models, including Augmented-{ESM1b ^22^, EVMutation ^68^, Evotuned UniRep ^69^, DeepSequence ^67^}, on the fitness data of 12 proteins from the original study ^27^. Using 240 randomly sampled mutants for training and the rest for testing, SPURS-augmented models outperformed their counterparts for most proteins, with a 7% improvement in Spearman correlation (Fig. 4b). To further assess performance, we extended evaluation to the ProteinGym benchmark ^29^, which includes over 200 deep mutational scanning (DMS) datasets spanning various protein fitness metrics such as catalytic activity, binding affinity, stability, and organismal fitness. We excluded DMS datasets measuring stability, so as to test generalization beyond the property SPURS was trained on. Across the resulting 141 DMS datasets, SPURS-augmented DeepSequence model outperformed the original Augmented DeepSequence–the most competitive method reported ^27^–in 115 (82%) cases, with an overall 15% improvement in Spearman correlation (Fig. 4c). Performance improvements were especially pronounced in datasets measuring expression (24.1%) and organismal fitness (16.5%) compared to activity (8.0%) and binding (12.6%) (Fig. 4d). These improvements were consistent across varying training set size *N* from as few as 48 to 240 variants, and up to 80% of the total sequences in a DMS dataset (Fig. 4e).

In summary, these results showed that the SPURS’s ΔΔ*G* predictions provide informative priors for protein fitness prediction, consistently enhancing leading low-*N* models across diverse DMS datasets and training data sizes. We note that our SPURS-augmented model itself is not intended to be a stand-alone, state-of-the-art predictor that outperforms a large volume of existing low-*N* models leveraging sophisticated deep learning models (e.g., pLMs) ^27,36,66^. Instead, it serves as a simple yet effective enhancement that yields consistent performance gains when added to already-competitive models, highlighting SPURS’s utility as a general-purpose stability prior for protein engineering tasks.

### SPURS reveals contribution of stability to pathogenicity

The human proteome harbors millions of missense variants, where single amino acid substitutions may alter protein structure and function, contributing to genetic diseases ^70–72^. While pathogenic variants may disrupt function through various molecular mechanisms such as perturbing interactions, the loss of function due to mutation effect on stability has been recognized as a major cause of disease ^73–77^. Given SPURS’s ability to predict protein stability changes (ΔΔ*G*) upon mutations, we sought to apply SPURS to examine the role of protein stability in pathogenicity.

We used SPURS to predict ΔΔ*G* for all possible single amino acid substitutions in the human proteome (178,987,201 variants across 19,652 proteins). Among these, 696,736 variants spanning 16,997 proteins have clinical annotations from ClinVar ^78^ as either (likely) ‘benign’, (likely) ‘pathogenic’, or variants of uncertain significance (VUS). Additionally, we included 15,820 missense variants from gnomAD (allele frequency >0.05) as putative benign variants ^40,79,80^. Our analysis revealed that pathogenic variants are more destabilizing, whereas most benign variants exhibit minimal stability changes (Fig. 5a). For example, 68% pathogenic variants (7,761/26,090) were destabilizing (ΔΔ*G >* +0.5 kcal/mol ^30,53^), compared to only 19% of benign variants (9,497/50,763). These findings align with previous studies showing loss of stability as a key driver for loss of function for many disease variants ^7,40,74,81^.

**Figure 5:**
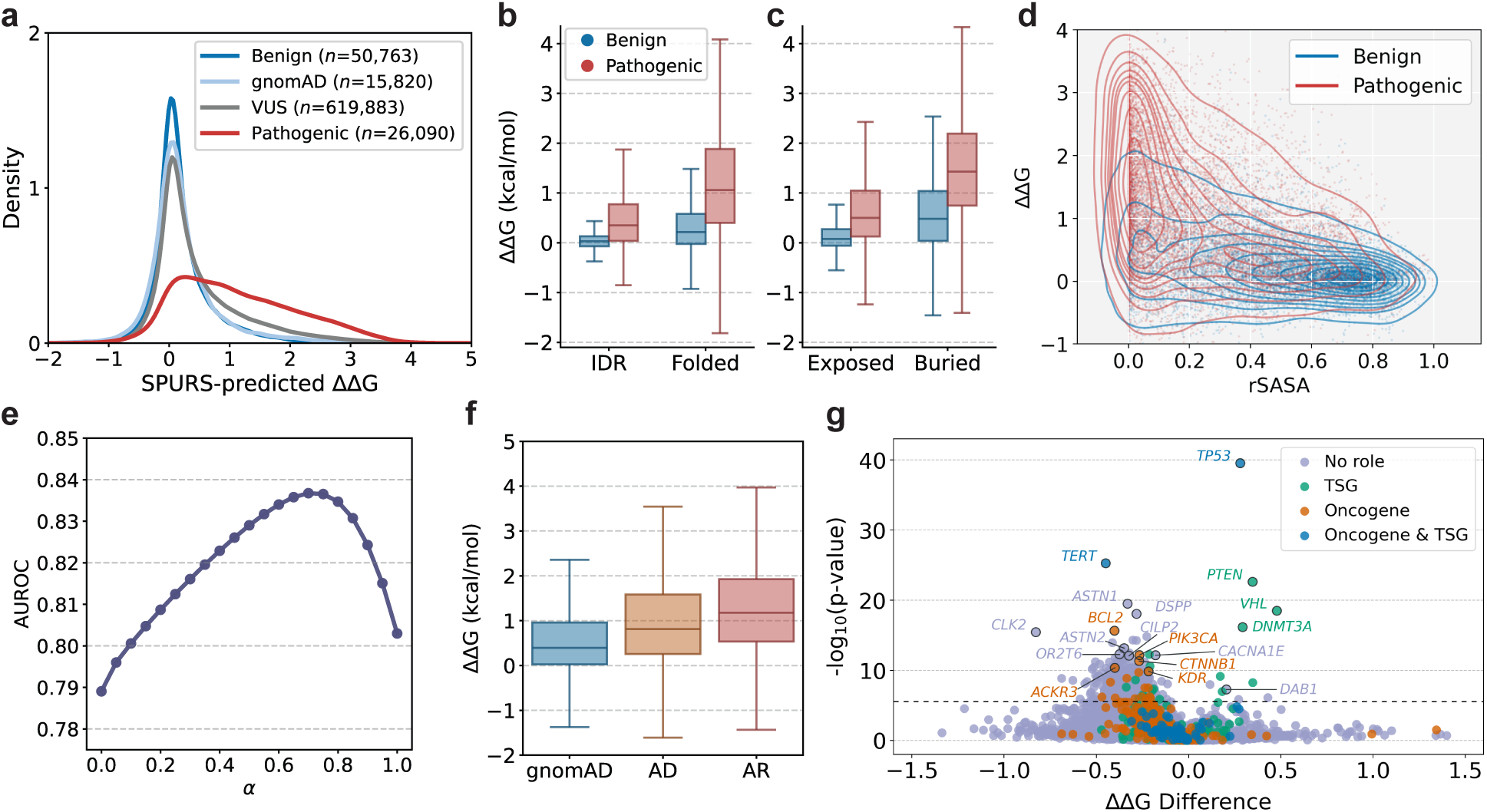
SPURS reveals the role of protein destabilization in variant pathogenicity. **a**, Distribution of SPURS-predicted stability change for ClinVar variants labeled as benign (including ‘Benign’ and ‘Likely benign’), pathogenic (including ‘Pathogenic’ and ‘Likely pathogenic’), and variants of uncertain significance (VUS). Common gnomAD variants (allele frequency >0.05) are included as putative benign variants for comparison. **b-c** Distribution of SPURS-predicted stability change for benign and pathogenic ClinVar missense mutations, categorized by (b) spatial locations—folded regions (*n*=31,463 benign, *n*=24,085 pathogenic) vs. intrinsically disordered regions (IDRs; *n*=19,207 benign, *n*=1,478 pathogenic)—and (c) solvent exposure—exposed residues (*n*=42,158 benign, *n* =10,419 pathogenic) vs. buried residues (*n* =8,512 benign, *n*=15,144 pathogenic). **d,** Joint distribution of SPURS-predicted ΔΔ*G* and relative solvent-accessible surface area (rSASA) for ClinVar benign and pathogenic missense variants. Contour maps indicate the density of variants in the rSASA–ΔΔ*G* space. **e,** AUROC for classifying ClinVar benign and pathogenic variants using a score that linearly interpolates SPURS-predicted ΔΔ*G* and rSASA, defined as *α·*ΔΔ*G*+(1*−α*)*·*rSASA, with *α* as the interpolation coefficient. **f,** Comparison of SPURS-predicted ΔΔ*G* values for gnomAD variants (*n*=15,820) and pathogenic ClinVar mutations in autosomal dominant (AD, *n*=6,722) and autosomal recessive (AR, *n*=5,603) genes. **g,** Gene-level enrichment of destabilizing or stabilizing cancer-associated missense mutations. ΔΔ*G* difference for a gene is defined as the mean SPURS-predicted ΔΔ*G* of observed COSMIC cancer mutations minus that of unobserved mutations in the same gene. A positive ΔΔ*G* difference indicates selection for destabilizing mutations in the gene, whereas a negative difference suggests selection against destabilization. The horizontal dashed line represents the Bonferroni-corrected significance threshold (*P* = 2.8 *×* 10*^−^*^6^, two-side Mann-Whitney U test), corresponding to a pre-correction significance cutoff of 0.05. Box plots in **b**, **c**, and **f** indicate the median (center line), 25th–75th percentiles (box), and whiskers represent data within 1.5*×* the interquartile range from the box.

To examine how structural context modulates the relationship between stability changes and pathogenicity, we stratified ClinVar variants by spatial locations and solvent exposure using AlphaFold2 predicted structures. First, we compared SPURS’s ΔΔ*G* predictions for mutations in folded regions versus intrinsically disordered regions (IDRs). In both groups, pathogenic variants were more destabilizing than benign ones (mean ΔΔ*G* difference +0.86 kcal/mol and +0.40 kcal/mol for folded regions and IDRs, respectively), but the effect was stronger in folded regions (Fig. 5b), consistent with previously reported tolerance of IDRs to mutations ^82^. While we recognize that AlphaFold2-predicted structures used in this analysis may not model IDRs accurately ^83^, recent studies on IDR conformation modeling may facilitate a better understanding of stability’s role in pathogenicity in IDRs ^84^.

Within folded regions, we compared ΔΔ*G* distributions for buried versus exposed residues (Fig. 5c). Again, pathogenic variants were more destabilizing than benign variants in both groups, but the difference was notably greater in buried positions than in exposed regions (mean ΔΔ*G* = 1.47 kcal/mol vs. = 0.67 kcal/mol). This observation, consistent with previous studies ^85,86^, suggests that pathogenic variants in buried residues primarily drive loss of function by reducing stability, whereas those at exposed sites likely disrupt function through alternative mechanisms, such as directly perturbing functional sites or intermolecular interactions ^86^.

Recognizing that the impact of stability on pathogenicity varies with residue exposure, we hypothesize that combining ΔΔ*G* prediction with structural context could enhance the identification of pathogenic mutations. Using relative solvent-accessible surface area (rSASA) to quantify residue exposure (Methods), we found that pathogenic variants were enriched in regions with both high ΔΔ*G* values and low rSASA scores (Fig. 5d), indicating that most pathogenic variants are located in buried residues and destabilize protein structure. Motivated by this, we developed an unsupervised predictor by linearly interpolating SPURS’s predicted ΔΔ*G* with rSASA with a weighting coefficient *α* to classify pathogenic and benign variants. Despite lacking supervised training on clinical variant annotations, this simple score achieved a maximum AUROC of 0.84 at *α* = 0.7, outperforming models based on either feature alone (Fig. 5e). This result highlights the potential synergy between stability and structural context in explaining pathogenicity. Note that our goal here is not to outperform state-of-the-art variant effect predictors with an optimal value of *α* ^54,87,88^, but rather to demonstrate that integrating stability prediction and structural context yields a synergistic and effective classifier. Together, these results (Figs. 5a-e) reinforce prior studies ^86,89^ and highlight the importance of structural contexts in understanding variant pathogenicity.

Next, we investigated the mutation effects on stability in different modes of inheritance and disease mechanisms. Autosomal recessive (AR) diseases are strongly associated with loss-of-function mutations, whereas autosomal dominant (AD) diseases can arise from alternate molecular mechanisms, including gain-of-function mutations, dominant-negative effect, or protein aggregation ^79,90^, which may not always involve substantial stability loss. We collected 6,722 AR-associated genes and 5,603 AD-associated genes annotated in the OMIM databases ^91^ and applied SPURS to predict ΔΔ*G* for the 12,325 pathogenic ClinVar variants covered by these genes. Consistent with previous findings ^40,79^, we found that pathogenic variants in AR genes were more destabilizing than those in AD genes or gnomAD variants (Fig. 5f; one-sided Mann-Whitney U test both *P <* 10^−50^). This difference is partly explained by the spatial distribution of mutations: as shown in prior studies ^92–95^ and our analysis (Fig. 5c), pathogenic mutations are more frequent in buried residues, where they have stronger destabilizing effects. Interestingly, it has been observed that AR mutations are particularly enriched in buried regions compared to AD and gnomAD variants ^79^. These findings highlighted the utility of SPURS with respect to disease mutations in understanding molecular mechanisms, implying that while stability loss is a major contributor to pathogenicity, a modest ΔΔ*G* effect does not necessarily indicate benignity, as certain pathogenic mutations may act through alternative mechanisms.

Finally, we analyzed stability effects in cancer-associated mutations. Genes that harbor cancer-driving mutations are broadly categorized into oncogenes, which promote cell proliferation, and tumor suppressor genes (TSGs), which inhibit uncontrolled cell growth. Given that tumorigenesis is often driven by gain-of-function mutations in oncogenes and loss-of-function mutations in TSGs ^96,97^, we leveraged SPURS’s ΔΔ*G* predictions to examine the molecular mechanisms of cancer-associated mutations. We obtained 5,225,814 somatic missense mutations observed across various cancer types from the COSMIC Cancer Mutation Census (CMC) database ^98,99^ (Methods). Within each gene, we compared SPURS’s predicted ΔΔ*G* for cancer mutations cataloged in COSMIC with that for all other possible missense mutations in the same gene (Fig.5g), following prior work ^97^. A gene with a positive ΔΔ*G* difference indicates that there has been selection for structurally damaging mutations in that gene (right side of Fig. 5g). Notably, this category was enriched for well-known TSGs, including *TP53*, *PTEN*, and *VHL*, suggesting that mutations in these genes likely drive tumorigenesis through stability loss. Conversely, genes with negative ΔΔ*G* difference (left side of Fig. 5g) were enriched for known oncogenes, such as *TERT*, *ACKR3*, *PIK3CA*, and *BCL2*. Several genes in this category, while not explicitly annotated in COSMIC, have literature support for tumorigenic roles, including *DSPP* ^100^ and *CLK2* ^101^. These results suggested that oncogenic mutations tend to avoid destabilization, likely favoring gain-of-function effects or disrupting intermolecular interactions rather than affecting structure stability. Overall, our analysis recapitulated previous findings ^96,97^, highlighting that TSGs mutations often lead to destabilization and loss of function, while oncogene mutations tend to exhibit selection for gain-of-function effects, with modest perturbation on stability.

Taken together, these findings illustrate SPURS’s capacity to discern the nuanced role of stability and structural context in variant pathogenicity, offering insights into the molecular mechanisms underlying human diseases.

## Discussion

We presented SPURS, a deep learning framework for protein stability prediction. One notable advantage of SPURS is its strong generalizability across proteins, stability measures (ΔΔ*G* or Δ*T_m_*), and datasets from different studies. This generalizability stems from two key design choices. First, SPURS is trained on Megas-cale, a high-coverage dataset encompassing all single and selected double mutants across hundreds of diverse proteins. Equally important is the algorithmic design in SPURS’s neural network architecture to fully harness the comprehensive stability measurements in Megascale. SPURS effectively integrates pre-trained sequence-and structure-based protein generative models (ESM2 and ProteinMPNN) through a neural network rewiring strategy, enabling them to be fine-tuned jointly for stability prediction in a data-efficient and scalable manner.

Our approach was motivated by previous findings that pre-trained protein generative models, when used in an unsupervised setting, already correlate well with experimentally measured ΔΔ*G* ^28–30,33^. We thus reasoned that adapting these pre-trained models using stability-specific fine-tuning would lead to stronger stability models. SPURS’s core innovation lies in the explicit integration of ESM2 and ProteinMPNN using a lightweight neural rewiring strategy. It integrates protein representations obtained from two different data modalities, fusing the structure embeddings that encode the geometric features important to protein stability with sequence embeddings that capture the evolutionary constraints in protein sequences. In addition, the resulting architecture is both expressive and parameter-efficient for fine-tuning while avoiding overfitting. While multimodal fusion has been well explored in fields such as vision-language modeling ^102^, and has recently inspired inverse folding studies ^42,103,104^, SPURS, to our knowledge, is the first to successfully integrate multi-modal protein generative models for stability prediction.

This study also addresses several limitations in prior stability prediction evaluations. First, in contrast to many previous studies that predominantly assessed models on random or in-distribution test splits, we systematically benchmarked SPURS on over ten independent datasets representing diverse proteins and various mutant coverage, where SPURS consistently outperformed state-of-the-art methods, highlighting its superior accuracy and generalizability. Furthermore, while most existing methods focus on single-mutation variants ^105^, we specifically evaluated SPURS’s ability to predict stability effects of higher-order variants, which suggested that SPURS effectively models non-additive mutational effects, outperforming additive baselines and models explicitly trained to capture epistasis. Importantly, we also addressed a common pitfall in model evaluation: the overreliance on correlation-based metrics, which are biased toward destabilizing mutations due to their overrepresentation ^50^. Instead, we reported classification-based performance in identifying stabilizing mutations–critical for protein engineering and design–where SPURS demonstrated higher precision and recall than leading method such as ThermoMPNN.

Another practical advantage of SPURS is its computational efficiency. By leveraging representation-sharing, it predicts ΔΔ*G* for all single mutations of a protein in only one forward pass. For high-order mutations, only a lightweight epistasis decoder is used, incurring minimal additional cost. This enables proteome-scale stability analysis: for instance, SPURS can perform site-saturation mutagenesis for the human proteome (19,652 proteins) in just 30 minutes on a single GPU.

Beyond stability prediction, we demonstrated the versatility of SPURS in various analyses in structural biology, protein engineering, and functional genomics. We showcased broad applications in which SPURS’s ΔΔ*G* predictions were used to identify functional residues in proteins, enhance protein fitness prediction, and investigate the contribution of stability change to pathogenicity in human genetic diseases. These applications demonstrate that SPURS’s superior accuracy, generalizability, and computational efficiency can serve as foundational capabilities to support diverse tasks in protein informatics.

Looking forward, we envision SPURS as a generalizable pre-trained model to predict stability changes for unseen proteins and mutations. With its state-of-the-art prediction accuracy, SPURS is a robust alternative to computationally expensive biophysical models for protein stability analysis. Since SPURS is trained on ΔΔ*G* data, its predictions for other stability measures, such as melting temperature (Δ*T_m_*), capture relative rankings rather than absolute values. A promising future direction is to exploit SPURS’s transferability to predict absolute magnitudes of other thermodynamic stability metrics, including Δ*T_m_* or Δ*G*, through dataset-specific fine-tuning. SPURS’s stability change predictions can also be combined with existing general variant effect predictors ^31,54,106–108^ and competitive models ^109–111^ in the Critical Assessment of Genome Interpretation (CAGI) ^112^ challenges to improve variant interpretation. SPURS also opens up exciting directions for protein engineering, either as a surrogate model for directed evolution or a reward model to guide generative AI models for designing stability-enhanced proteins. These broad applications establish SPURS as a versatile tool to enable faster and scalable analyses for characterizing protein stability and accelerating protein engineering for important applications in therapeutic design and synthetic biology.

## Methods

### Representations of input sequence and structure

SPURS takes as input a protein’s wild-type sequence **x** = (*x*_1_*, …, x_L_*), where *L* is the sequence length, *x_i_* ∈ A denotes the amino acid at position *i* from the set A of 20 canonical amino acids, and predicts thermostability change (ΔΔ*G*) for all possible single-mutation variants and specified higher-order variants (Fig. 1). When available, SPURS also incorporates the wild-type protein’s 3D structure. If no experimentally determined structure is accessible, SPURS uses AlphaFold2^5^ to predict the structure. In our experiments, experimental structures were used when the original benchmark datasets (see ‘Datasets’) provided a Protein Data Bank (PDB) ID for the protein; otherwise, AlphaFold-predicted structures were used, following the practice of previous studies on stability prediction ^29,38^. The structure source for each benchmark dataset is detailed in Supplementary Table 6.

SPURS integrates ProteinMPNN ^26^ as a structure encoder and ESM2^24^ as both a sequence encoder and mutation effect predictor. ProteinMPNN is a message-passing neural network (MPNN) ^113,114^, a type of graph neural network, which learns representations from a 3D protein structure. Structure information is described as atomic coordinates S = {***c****_i_* ∈ R*^Nb^* ^×3^}*^L^*, where ***c****_i_* contains the coordinates of *N_b_* backbone atoms (e.g., *C*, *C_α_*, *O*, and *N* atoms) for residue *i*. This structure is abstracted into a graph with atoms as nodes and edges formed by connecting each atom to its 48 nearest neighbors. This structure graph is passed through three encoder and three decoder layers in the MPNN to yield per-residue embeddings in R*^L^*^×128^. To improve representation learning, we replaced the autoregressive decoding in the original ProteinMPNN model with a masked decoding scheme (described below). We concatenated MPNN’s output embedding with its internal residue embeddings (R*^L^*^×128^) learned based on the amino acid identity, resulting in combined embeddings ***z****_m_* ∈ R*^L^*^×2^^56^, which were then up-sized by a linear layer to ***z****_s_* ∈ R*^L^*^×1280^, referred to as structure features. In parallel, ESM2, a 33-layer Transformer ^1^^15^, encodes the protein sequence as a per-residue embedding ***z****_e_* ∈ R*^L^*^×1280^. Since ESM2 is pre-trained on massive natural sequence data, the output embeddings ***z****_e_* capture evolutionary patterns of protein language, referred to as evolutionary features.

### Model rewiring to learn structure-enhanced evolutionary features

To integrate the structure features ***z****_s_* ∈ R*^L^*^×1280^ captured by ProteinMPNN into the evolutionary features ***z****_e_* ∈ R*^L^*^×12^^80^ learned by ESM2, SPURS introduces a parameter-efficient rewiring mechanism based on Adapter layers ^43^, which has proven effective in fine-tuning large language models and for protein sequence design ^42,116^. Injecting structure-derived context into the sequence-based representation, the Adapter creates structure-enhanced evolutionary features ***z****_a_* ∈ R*^L^*^×12^^80^ by cross-attention followed by residual learning and nonlinear transformation:

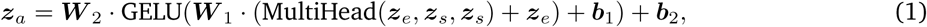

where MultiHead(*Q, K, V*) is the multi-head attention layer ^115^, with query *Q*, key *K*, and value *V* matrices. Here, we used ***z****_e_* as the query and ***z****_s_*as both key and value, allowing evolutionary features to attend to and create interactions with structural contexts. The attention output is added back to ***z****_e_* through a residual connection ^117^, followed by a two-layer MLP parameterized by ***W*** _{1,2}_ and ***b***_{1,2}_, with a GELU activation function ^118^, to produce the structure-enhanced evolutionary features ***z****_a_*. By learning to extract and integrate features from both structural and evolutionary contexts, SPURS is expected to provide more informed stability prediction with the enhanced feature ***z****_a_*. The Adapter is inserted after the 31st layer (out of 33 layers) of ESM2, with the position selected via hyperparameter search on the validation set. After the Adapter, ***z****_a_* is passed through the final layers of ESM2 and projected from R*^L^*^×12^^80^ to R*^L^*^×1^^28^ via a linear layer. Finally, we concatenate this projected output with the earlier ProteinMPNN embeddings ***z****_m_*∈ R*^L^*^×256^ to obtain the final protein embedding ***z****_o_* ∈ R*^L^*^×384^, reinforcing the structure prior while preserving the evolutionary context.

Building on prior work showing that Adapter-only fine-tuning can achieve comparable performance to fine-tuning the full Transformer model ^42,43^, we froze the ESM2 parameters (650 million parameters) and optimized only the Adapter and ProteinMPNN parameters (9.9 million parameters). This strategy (Fig. 1) reduced the number of trainable parameters by 98.5% compared to updating the full SPURS model, without compromising prediction accuracy.

### Efficient stability prediction module

To predict the stability changes, the embedding ***z****_o_* is passed to a multi-layer perceptron (MLP) *g*: R^3^^84^ → R^20^, with each output dimension corresponds to one of the 20 amino acids (AAs). This MLP, shared across all *L* positions in the sequence, projects ***z****_o_* ∈ R*^L^*^×384^ to a matrix ***ϕ*** = *g*(***z****_o_*) ∈ R*^L^*^×20^. The element of this *L* × 20 matrix ***ϕ*** is indexed by the sequence position and AA type, in which ***ϕ***(*i, a*) represents a trainable weight that approximates the thermostability (ΔG) when the amino acid at residue *i* is of type *a*.

This matrix ***ϕ*** can be used to derive the change in Δ*G* (i.e., ΔΔ*G*) for a single mutation. Denote ***x***^WT^ and ***x***^MT^ as the wild-type sequence and its single-mutation variant resulting from the substitution *x_i_*^WT^ → *x_i_*^MT^ at position *i*. The stability change of ***x***^MT^ with respect to ***x***^WT^ is defined as

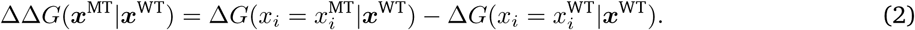

In analogy, SPURS (denoted as *f_θ_* parameterized by *θ*) predicts the ΔΔ*G* for the variant ***x***^MT^ with a substitution at residue *i*, conditioned on the protein’s wild-type sequence ***x***^WT^ and structure S^WT^, as following:

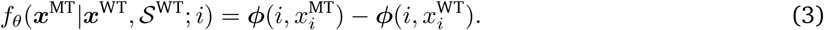

This formulation enables efficient and scalable ΔΔ*G* prediction: the matrix ***ϕ*** only needs to be computed once in a single forward pass of the neural network, and then can be reused efficiently to derive the ΔΔ*G* for all single mutations using Eq. 3. In contrast, many existing methods use mutant sequences as input, which requires O(*L* × 20) forward passes to predict ΔΔ*G* for all *L* × 20 single-mutation variants. By conditioning on the wild-type sequence and structure, SPURS predicts all single-mutation variants altogether in O(1) pass, significantly improving prediction efficiency. This approach, also adopted in recent studies ^36,51^, is well-suited for large-scale protein stability analysis.

SPURS was trained using a mean squared error (MSE) loss to minimize the difference between predicted and experimentally measured ΔΔ*G* values. Each training data batch includes all mutants derived from a single wild-type sequence. Training was performed on an NVIDIA A40 GPU for a maximum of 200 epochs using the AdamW optimizer ^119^ with a learning rate of 0.0001. A plateau scheduler was used for adaptive learning rate adjustment, and early stopping was employed to prevent overfitting by terminating training once the validation performance was not improved for 30 epochs. Hyperparameters such as batch size, learning rate, and optimizer settings were selected using the Megascale validation set.

### Extension to higher-order mutations

The prediction scheme described above for single mutations can be extended to higher-order variants. Given a mutant sequence ***x***^MT^ of length *L* that differ from the wild-type sequence ***x***^WT^ at multiple positions, we define the set of mutated positions as M = {*i*|*x_i_*^MT^ ≠ *x_i_*^WT^; 1 ≤ *i* ≤ *L*}. The stability change induced by multiple mutations is often not simply the additive effects of individual point mutations due to epistatic interactions between residues. SPURS thus predicts the ΔΔ*G* for ***x***^MT^ as

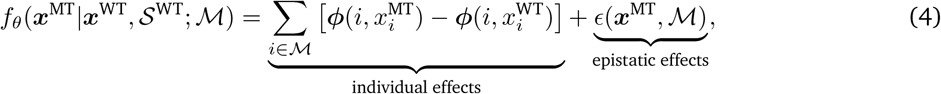

where the first term aggregates individual mutation effects derived from previously computed matrix *ϕ*, and the epistatic effect *ɛ* among mutated residues is added to predict the overall ΔΔ*G*. The epistatic term *ɛ* is predicted by a dedicated MLP: *ɛ*(***x***^MT^, M) = *g_ɛ_*(***z***^M^), where *g_ɛ_* is a four-layer MLP network with hidden dimensions 768, and ***z***^M^ is a latent mutation representation that encodes features of all mutations in M. We factorized the representation ***z***^M^ into the representations of its compositional point mutations: ***z***^M^ =P***z***^(*i*)^. The representation of a point mutation ***z***^(*i*)^ is decomposed as the concatenation of a context-dependent feature and a context-independent feature: ***z***^(*i*)^ = ^(^***z***^(*i*)^∥:***e*** (*x*^MT^)^i^ ∈ R^512^, where ∥: is the vector concatenation, ***z***^(*i*)^ ∈ R^384^ is a context-dependent feature extracted from the *i*-th row of the aforementioned per-residue embedding matrix ***z****_o_*∈ R*^L^*^×384^, whereas ***e***_AA_(*a*): A → R^128^ is an embedding layer that maps an amino acid *a* ∈ A to a context-independent feature in R^128^. Intuitively, ***z***^(*i*)^ captures the contextualized features of position *i* in the wild-type sequence, and ***e***_AA_(*x*^MT^) informs the mutated amino acid type.

The formulation in Eq. 4 allows efficient and scalable ΔΔ*G* prediction for high-order mutations. Similar to the prediction of single-mutation variants, SPURS only requires one forward pass of the rewired ESM and ProteinMPNN networks to compute the per-residue embeddings ***z****_o_* and matrix *ϕ*, which can be reused to infer ΔΔ*G* for all possible single or higher-order variants. For each specific combination of mutations, SPURS only needs to run one additional forward pass on *g_ɛ_*, adding only minimal computational overhead (less than 0.1% of the total parameters of ESM2 and ProteinMPNN).

To train this extended architecture of SPURS, we randomly initialized the parameters of *g_ε_* and retained the weights of all other layers optimized from the training on Megascale single-mutation data. We then fine-tuned the entire model using 122,278 double-mutation variants from Megascale training set. Double-mutation variants from the Megascale validation set were used for hyperparameter tuning, and those in the test split were used to evaluate SPURS’s predictive performance on higher-order mutations (Fig. 2f).

### Improved ProteinMPNN decoding for structural feature learning

ProteinMPNN was originally designed as an inverse folding model to generate sequences compatible with a given backbone structure. It uses an autoregressive (sequential) decoding scheme to generate each amino acid one at a time, conditioning on the input structure and already decoded residues: *p*(*x_i_*|***x****_<i_*; S) where ***x****_<i_* = *x*_1_ *… x_i_*_−1_. While this design aligns with ProteinMPNN’s initial purpose, it inherently limits its ability to fully learn structural features for stability prediction in our work, as the model cannot access information from the entire sequence during decoding.

To overcome this limitation, we adopted a one-shot decoding strategy, allowing each residue in the sequence to access information from all other positions simultaneously: *p*(*x_i_*|***x***_−*i*_; S) where ***x***_−*i*_ denotes the full sequence excluding residue *i*, thereby providing richer sequence context and improving structure representation learning. To fully leverage the advantages of this new strategy, we fine-tuned the ProteinMPNN parameters jointly with the Adapter and prediction modules during SPURS’s training, rather than keeping them fixed, allowing the model to better adapt to the refined decoding scheme and contribute more informative structure features for stability prediction.

### Functional sites identification

For a given wild-type protein sequence, we used SPURS to predict ΔΔ*G* and ESM1v ^28^ to predict the sequence likelihood change for all possible single-residue substitutions. The ESM-predicted sequence likelihood change, or delta log-likelihood (ΔLL), is defined as the difference in log-probability between a mutant and wild type residue at position *i*:

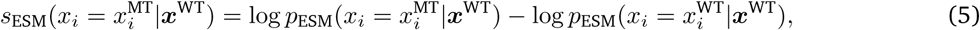

where *p*_ESM_(*x_i_*= *a*|***x***) is the ESM-predicted probability of amino acid *a* occurring at position *i*, given the wild-type sequence ***x*** as context ^28^. The score *s*_ESM_(*x_i_* = *x*^MT^|***x***^WT^) can be interpreted as the relative evolutionary fitness of mutant ***x***^MT^, which has been shown to be an effective zero-shot predictor of experimentally measured fitness data ^28,29^.

Inspired by prior studies modeling the relationship between the free energy changes due to mutations in protein folding and binding using a non-linear Boltzmann distribution ^56,59–62^, we applied a sigmoid function to fit the non-linear relationship between a mutant’s stability change and evolutionary fitness. Specifically, we defined a stability change score *s*_stab_(*x_i_*= *x*^MT^|***x***^WT^) as the negative SPURS-predicted ΔΔ*G* (i.e., −*f_θ_*(·)) and min-max normalized this score across all single-mutation variants, with the minimum set to the 0.1% percentile and the maximum to the 99.9% percentile. In this way, stabilizing mutations are assigned *s*_stab_ scores close to 1, while destabilizing ones approach 0.

Next, we fit a sigmoid function on the *s*_stab_ and *s*_ESM_ scores of all single-mutation variants for each protein:

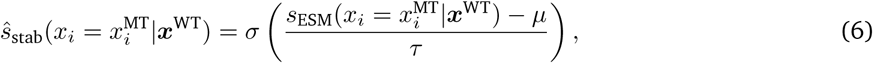

where *σ*(*z*) = 1*/*(1 + *e*^−*z*^) is the standard sigmoid function, *µ* and *τ* are learnable parameters controlling the sigmoid curve’s shape (location and steepness), and *s*^_stab_ is the sigmoid-fit values of *s*_stab_. To prioritize fitting the low-stability variants, we weighted a variant by SPURS’s ΔΔ*G* prediction, following the Domainome study ^40^:

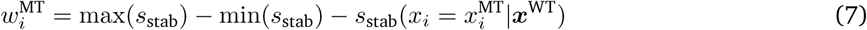

The residue of the fit is defined as:

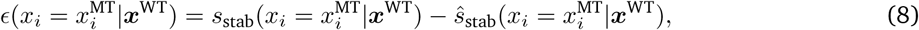

which reflects how much the predicted stability deviates from what would be expected based on the predicted fitness alone. The Domainome study showed that these residuals may indicate mutations with larger or smaller effects on fitness than can be accounted for by changes in stability ^40^, suggesting stronger functional constraints at site *i*. We thus define a per-site importance score as the average residuals across the 20 AA mutations at a site, referred to as function score:

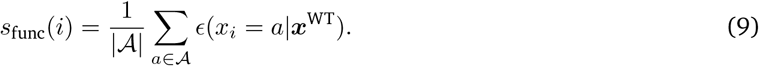

As shown in the Domainome study ^40^ and confirmed by our results (Fig. 3), a larger *s*_func_(*i*) value suggests that site *i* is more likely a functional site.

### Enhanced protein fitness prediction with SPURS

To demonstrate that SPURS can enhance supervised prediction of general protein fitness, we used the “Aug-mented models” developed by Hsu et al.^27^, a leading low-*N* fitness prediction method, as our base model. To predict the quantitative fitness value of a variant, Augmented models represent the input sequence as a concatenation of one-hot encoding of the amino acids in its sequence, resulting in a flattened vector of dimension *L* ×|A|, where A is the set of 20 canonical amino acids. This vector is augmented to a (*L* ×|A|+ 1)-dimension feature vector by incorporating a scalar representing the input sequence’s evolutionary plausibility (the delta log-likelihood score) predicted by a protein sequence generative model, such as a pLM (e.g., *s*_ESM_ in Eq. 5) or other sequence likelihood models (e.g., EVmutation ^120^, Evotuned UniRep ^69^, DeepSequence ^67^). A Ridge regression model is then trained on these augmented features to predict experimental fitness measurements.

We extended this approach by incorporating the ΔΔ*G* value predicted by SPURS as an additional input feature, resulting in a (*L* × |A| + 2)-dimensional feature vector. We refer to this enhanced model as the SPURS-augmented model. To benchmark performance, we followed the experimental design of Hsu et al.^27^. For each deep mutational scanning (DMS) dataset, we first set aside 20% randomly sampled variants as the test set. From the remaining variants, we sampled *N* mutants as the training set, where *N* was set to 48, 96, 144, 192, 240, or *N* = 80% of total sequences in a DMS dataset, following the setup from Hsu et al. ^27^. For each value of *N*, we repeated the sampling and model training 20 times with different random seeds and reported the mean performance across repetitions as the final evaluation metric.

### Analysis of stability effects on pathogenicity

We obtained the missense variants and their clinical annotations from the ClinVar database ^78^, following the processing protocol in Cagiada et al.^86^. In our analysis, we only included variants annotated as ‘benign’, ‘likely benign’, ‘pathogenic’, ‘likely pathogenic’, or ‘variants of uncertain significance’ (VUS). In our analysis, variants with ‘benign’ or ‘likely benign’ annotations were grouped as benign, and those with ‘pathogenic’ or ‘likely pathogenic’ annotations were grouped as pathogenic. We applied SPURS to predict ΔΔ*G* for all possible missense mutations across the human proteome (19,652 proteins), and 696,736 variants from 16,997 proteins were successfully mapped to ClinVar annotations.

AlphaFold2 (AF2)-predicted structures of the mapped human proteins and their associated residue-level predicted Local Distance Difference Test (pLDDT) scores were retrieved from the AlphaFold Protein Structure Database ^121^. Residues were categorized as folded if pLDDT > 50. Solvent exposure was estimated using relative solvent-accessible surface area (rSASA), defined as the SASA of a residue divided by its maximum SASA in a Gly-X-Gly tripeptide configuration. rSASA values were computed using the Biotite package ^122^ with normalization constants from Tien et al.^123^; residues with rSASA > 0.2 were considered exposed.

To jointly model the impact of structure and stability on pathogenicity, we interpolated ΔΔ*G* and rSASA scores after clipping them to the 0.1st–99.9th percentile range to reduce the influence of outliers. These values were then min-max normalized to the [0,1] interval, preserving relative differences while ensuring scale compatibility across the dataset.

The disease inheritance annotations (autosomal dominant [AD] or autosomal recessive [AR]) for genes were obtained from the OMIM database ^91^, following the approach of Gerasimavicius et al.^79^. We additionally included 15,820 gnomAD (v4.1) missense variants with allele frequency >0.05 as a reference set of putative benign variants. Cancer-associated mutations were collected from the Cancer Mutation Census dataset ^99^ (v99, GRCh38) in the COSMIC database ^98^. For each gene, a ΔΔ*G* difference score was computed as the mean predicted ΔΔ*G* of observed cancer-associated mutations minus that of all remaining possible missense mutations in that gene. The significance of pairwise comparison was assessed using the Mann-Whitney U test, with Bonferroni correction for multiple testing. The threshold of the corrected *P*-value was defined as *α/N*, where *α* = 0.05 is the commonly used significance level, and *N* = 18, 073 is the number of genes considered.

### Datasets

We used the training, validation, and test splits of the Megascale dataset created by Dieckhaus et al.^38^, which ensured no sequences sharing *>* 25% identity across splits. For benchmarking, we additionally collected ten independent test sets, including Fireprot (HF) ^18,38^, Ssym-direct ^47^, Ssym-inverse ^47^, S669^9^, S783^8^, S2648^15^, S461^124^, S8754^45^, S4346^45^, S571^45^. Among these, four datasets (Fireprot, Ssym-direct, Ssym-inverse, and S669) were pre-filtered and provided by Dieckhaus et al.^38^, while the remaining six datasets were obtained from Xu et al.^45^. Fireprot (HF) is a homology-free subset of the Fireprot dataset, curated to ensure that sequences share less than 25% identity with those in Megascale. We used experimentally solved structures from the PDB database to run SPURS and structure-based baseline methods if the representative structures of the wild-type proteins were provided by the original studies of these test sets; otherwise, AlphaFold2-predicted structures were used.

We first trained SPURS using single-mutation data in Megascale training and validation splits and evaluated it using the Megascale test split and the ten independent test sets (statistics provided in Supplementary Table 6). To prevent data leakage across evaluation sets, any training sequences with >25% sequence identity to sequences in the independent test sets were removed using MMseqs2^125^. To enable predictions for higher-order mutations, double-mutant variants from the Megascale training and validation splits (statistics provided in Supplementary Table 7) were incorporated into training for the extended model with the epistasis module *g_ɛ_*. Double mutants in the Megascale test split were used to assess the performance of this extended model.

Functional site annotations for Domainome sequences were retrieved from the Conserved Domain Database (CDD) by parsing the National Center for Biotechnology Information (NCBI) protein entry page (e.g., https://www.ncbi.nlm.nih.gov/protein/UNIPROT_ID/, replacing “UNIPROT_ID” with the corresponding UniProt identifier). For the enzyme in Fig.3 (Alcohol dehydrogenase, Uniprot ID: P00327, PDB ID: 1QLH), functional site annotations were obtained from the Mechanism and Catalytic Site Atlas (M-CSA; https://www.ebi.ac.uk/thornton-srv/m-csa/). Experimental fitness measurements and baseline predictions for proteins in Fig.4 were taken from the ProteinGym benchmark ^29^ (https://proteingym.org/).

## Data availability

Unless otherwise stated, all data supporting the results of this study can be found in the article, supplementary, and source data files. We used *esm2_t33_650M_UR50D* checkpoint of ESM2 (https://dl.fbaipublicfiles.com/fairesm/models/esm2_t33_650M_UR50D.pt), *v_48_020* checkpoint of ProteinMPNN (https://github.com/dauparas/ProteinMPNN/blob/main/vanilla_model_weights/v_48_020.pt), and *esm1v_t33_650M_UR90S_1* checkpoint of ESM1v (https://dl.fbaipublicfiles.com/fairesm/models/esm1v_t33_650M_UR90S_1.pt). The Megascale dataset was downloaded from https://zenodo.org/records/7844779. Megascale split, Fireprot(HF), S669, Ssym-direct, and Ssym-inverse were downloaded from https://github.com/Kuhlman-Lab/ThermoMPNN. S783, S2648, S461, S8754, S4346 and S571 were downloaded from https://github.com/Gonglab-THU/GeoStab. Domainome data and the performance of baseline models were downloaded from https://zenodo.org/records/11260616. Clinvar annotations were collected from https://ftp.ncbi.nlm.nih.gov/pub/clinvar/tab_delimited/ ^86^. Cancer-associated mutations were obtained from the Cancer Mutation Census v99 (GRCh38) dataset, specifically the file CancerMutationCensus_AllData_Tsv_v99_GRCh38, available at https://cancer.sanger.ac.uk/cosmic/download/cancer-mutation-census/v99/alldata-cmc. Source Data are provided with this paper.

## Code availability

SPURS was implemented in Python using the PyTorch (v1.12.0) library. The source code is available at https://github.com/luo-group/SPURS.

## Acknowledgements

This work is supported in part by the National Institute of General Medical Sciences of the National Institutes of Health under award R35GM150890 (Y.L.). The authors acknowledge the computational resources provided by the Delta GPU Supercomputer at NCSA of UIUC through allocation CIS230097 (Y.L.) from the Advanced Cyberinfrastructure Coordination Ecosystem: Services & Support (ACCESS) program, which is supported by NSF grants #2138259, #2138286, #2138307, #2137603, and #2138296 (Y.L.), and the cloud computational resources provided by Microsoft Azure through the Cloud Hub program at GaTech IDEaS and the Microsoft Accelerate Foundation Models Research (AFMR) program (Y.L.).

## Author contributions

Y.L. conceived and supervised the project Z.L. developed the computational framework and performed the evaluation analyses. Y.L. and Z.L. wrote the manuscript.

## A Supplementary Tables

**Supplementary Table 1:** Spearman correlation of SPURS and baseline models on test sets. This table provides the raw correlation results underlying Fig. 2d. Fireprot (HF) denotes a homology-free subset of the Fireprot dataset, curated to ensure that sequences share less than 25% identity with those in Megascale, enabling rigorous evaluation of model generalization.

|  |  | ESM | ProteinMPNN | FT-ESM | Models<br>FT-ProteinMPNN | ThermoMPNN | ProteinMPNN + ESM | SPURS |
| --- | --- | --- | --- | --- | --- | --- | --- | --- |
| Model Components | ESM Encoder | ✓ |  | ✓ |  |  | ✓ | ✓ |
|  | ProteinMPNN Encoder |  | ✓ |  | ✓ | ✓ | ✓ | ✓ |
|  | Finetuning |  |  | ✓ | ✓ | ✓ | ✓ | ✓ |
|  | Rewiring |  |  |  |  |  |  | ✓ |
| Datasets | Megascale | 0.33 | 0.41 | 0.64 | 0.70 | 0.72 | 0.72 | **0.74** |
|  | Fireprot (HF) | 0.43 | 0.55 | 0.48 | 0.64 | 0.64 | 0.57 | **0.67** |
|  | Ssym-direct | 0.36 | 0.58 | 0.43 | 0.71 | 0.71 | 0.60 | **0.76** |
|  | Ssym-inverse | 0.38 | 0.38 | 0.28 | 0.63 | 0.59 | 0.45 | **0.68** |
|  | S669 | 0.37 | 0.43 | 0.46 | 0.46 | 0.44 | 0.45 | **0.50** |
|  | S461 | 0.29 | 0.50 | 0.65 | 0.60 | 0.61 | 0.65 | **0.66** |
|  | S783 | 0.30 | 0.52 | 0.59 | 0.69 | 0.70 | 0.64 | **0.71** |
|  | S8754 | 0.16 | 0.24 | 0.53 | 0.60 | 0.61 | 0.52 | **0.64** |
|  | S2648 | 0.18 | 0.31 | 0.53 | 0.65 | 0.65 | 0.61 | **0.69** |
|  | S571 | 0.27 | 0.43 | 0.40 | 0.42 | 0.39 | 0.39 | **0.47** |
|  | S4346 | 0.32 | 0.52 | 0.46 | 0.61 | 0.60 | 0.51 | **0.64** |
|  | Avg. | 0.31 | 0.44 | 0.50 | 0.61 | 0.61 | 0.56 | **0.65** |

**Supplementary Table 2:**
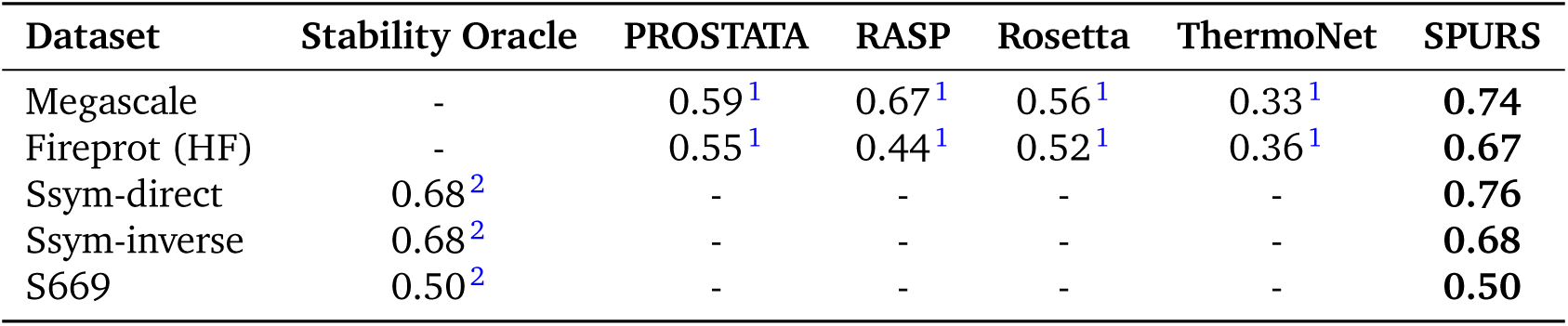
Spearman correlation of additional baseline models on test sets. Spearman correlation results for a group of additional baseline methods reported in Diaz et al. ^2^ and Dieckhaus et al. ^1^. These results were also presented in Fig. 2d.

**Supplementary Table 3:** Pearson correlation of SPURS and baseline models on test sets. Evaluation results following the same protocol as in Supplementary Table 1, using Pearson correlation as the performance metric to complement the Spearman results.

| Dataset | ESM | ProteinMPNN | FT-ESM | FT-ProteinMPNN | ThermoMPNN | SPURS |
| --- | --- | --- | --- | --- | --- | --- |
| Megascale | 0.34 | 0.49 | 0.68 | 0.74 | 0.75 | **0.78** |
| Fireprot (HF) | 0.33 | 0.48 | 0.48 | 0.63 | 0.64 | **0.66** |
| Ssym-direct | 0.30 | 0.59 | 0.44 | 0.71 | 0.70 | **0.74** |
| Ssym-inverse | 0.27 | 0.42 | 0.28 | 0.60 | 0.58 | **0.65** |
| S669 | 0.25 | 0.31 | 0.42 | 0.43 | 0.39 | **0.46** |
| S461 | 0.32 | 0.38 | **0.64** | 0.58 | 0.58 | **0.64** |
| S783 | 0.43 | 0.55 | 0.58 | 0.71 | 0.72 | **0.73** |
| S8754 | 0.37 | 0.38 | 0.47 | 0.57 | 0.58 | **0.60** |
| S2648 | 0.29 | 0.50 | 0.52 | 0.65 | 0.64 | **0.67** |
| S571 | 0.18 | 0.22 | 0.36 | 0.31 | 0.25 | **0.36** |
| S4346 | 0.27 | 0.43 | 0.44 | 0.54 | 0.53 | **0.58** |

**Supplementary Table 4:** Performance on double mutants in the Megascale test set. Evaluation of SPURS and baselines for predicting ΔΔ*G* values of higher-order (double) mutations. “Additive” predict ΔΔ*G* for high-order mutations by simply adding the predicted ΔΔ*G* values for constituent individual mutations. Results correspond to Fig. 2f.

| Model Name | Performance |
| --- | --- |
| DDGun-3D | 0.36 |
| DDGun-seq | 0.33 |
| ESM2 (Additive) | 0.40 |
| MPNN (Additive) | 0.16 |
| TheomMPNN (Additive) | 0.45 |
| SPURS (Additive) | 0.52 |
| MutateEverything | 0.54 |
| SPURS | 0.59 |

**Supplementary Table 5:**
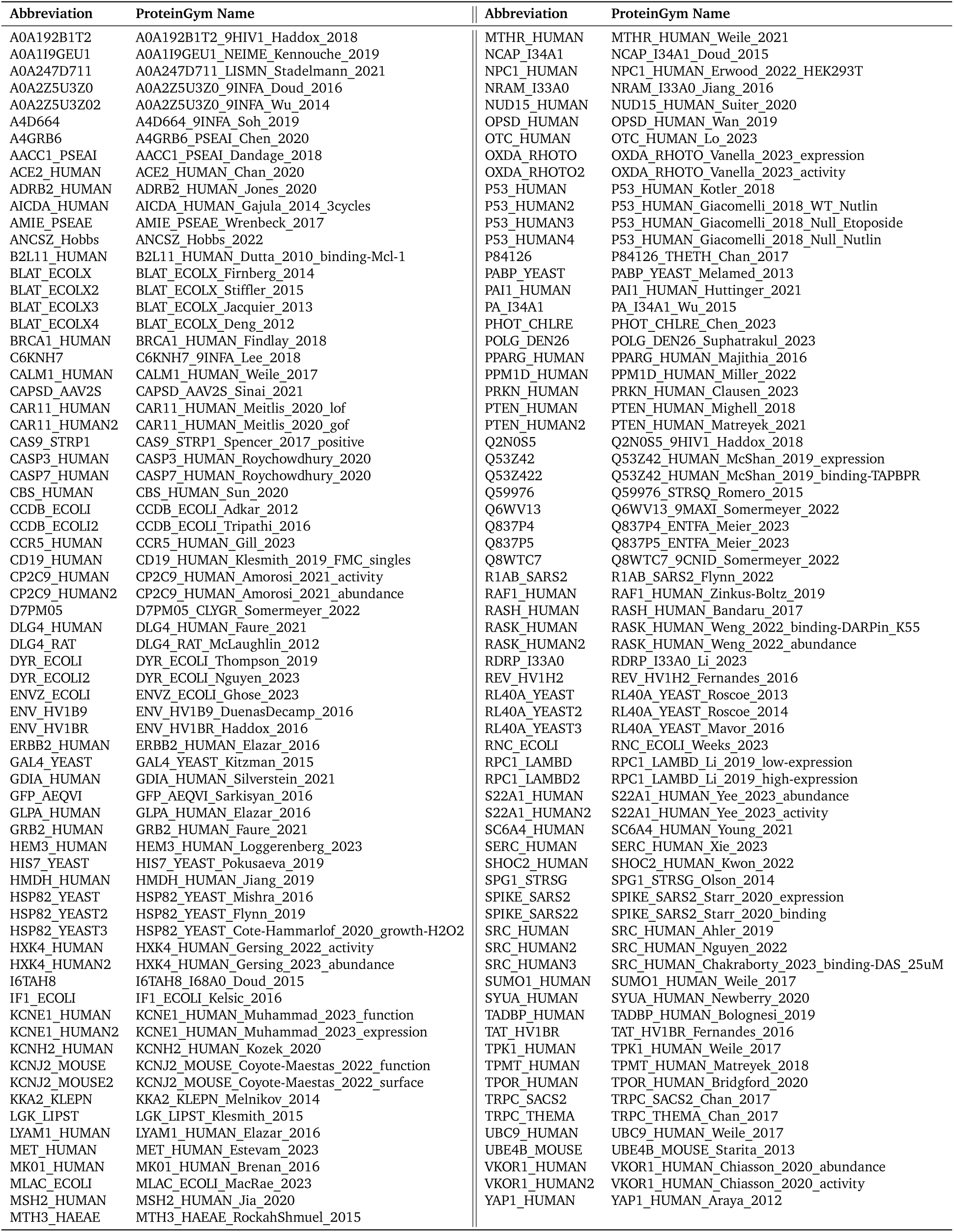
Abbreviations for DMS datasets. Digit suffixes distinguish multiple DMS studies on the same protein.

**Supplementary Table 6:** Summary statistics of single-mutation datasets used for training and evaluation. Number of proteins and total mutants per dataset. The “Structure” column denotes the source of structural data used in our experiments–AF2: AlphaFold2 predictions; PDB: experimentally solved structures available in the PDB database.

| Dataset | Number of mutants | Number of proteins | Structure |
| --- | --- | --- | --- |
| Megascale (training) | 216,919 | 239 | AF2 |
| Filtered Megascale (training) | 203,789 | 212 | AF2 |
| Megascale (validation) | 27,481 | 31 | AF2 |
| Filtered Megascale (validation) | 26,548 | 29 | AF2 |
| Megascale (test) | 28,312 | 28 | AF2 |
| Fireprot (HF) | 3,438 | 89 | PDB |
| Ssym-direct | 342 | 15 | PDB |
| Ssym-inverse | 342 | 342 | PDB |
| S669 | 669 | 94 | PDB |
| S461 | 461 | 48 | PDB |
| S783 | 783 | 55 | PDB |
| S8754 | 8,236 | 274 | PDB |
| S2648 | 2,648 | 131 | PDB |
| S4346 | 3,988 | 281 | PDB |
| S571 | 571 | 37 | PDB |
| Domainome | 536,164 | 522 | PDB |

**Supplementary Table 7:** Summary of double-mutation variants in Megascale. The number of proteins and double-mutation variants included in the Megascale training, validation, and test splits used for training and evaluating SPURS’s high-order mutation prediction module (Fig. 2f). The training/validation/test splits were obtained from the ThermoMPNN study ^1^.

| Dataset | Number of mutants | Number of proteins |
| --- | --- | --- |
| Megascale (training) | 122,278 | 116 |
| Megascale (validation) | 10,672 | 17 |
| Megascale (test) | 22,081 | 20 |

## B Supplementary Notes

### B.1 Baseline Methods

We compared SPURS against several leading stability predictors, including biophysics and machine learning models. For some baselines, we reused evaluation results reported in the original publications or in Dieckhaus et al.^1^, as noted below. It is worth noting that advantages were given to these baselines, as their training data are supersets of our filtered Megascale training set and may include proteins from the benchmark test sets used in our evaluation.

#### ThermoMPNN

Dieckhaus et al.^1^ developed ThermoMPNN, a transfer learning framework based on the ProteinMPNN architecture, trained on the Megascale dataset. The model uses a lightweight attention layer and MLP to transform ProteinMPNN embeddings into stability scores. We adopted the official implementation from https://github.com/Kuhlman-Lab/ThermoMPNN. Following the original setup, we first trained ThermoMPNN on the full Megascale dataset and reproduced the authors’ reported results on the Megascale test set, Fireprot (HF), and S669 benchmarks. For fair comparison, we then retrained the model using our filtered Megascale dataset and kept all original hyperparameters unchanged.

#### RASP

RASP, proposed by Blaabjerg et al.^3^, is a stability prediction model pretrained on a structural representation learning task and subsequently fine-tuned on Rosetta-predicted ΔΔ*G* values. We compared against the results reported by Dieckhaus et al.^1^.

#### Stability Oracle

Stability Oracle is a structure-based ΔΔ*G* predictor introduced by Diaz et al.^2^, based on a graph transformer architecture. It models the local structural environment surrounding a mutation site and predicts position-specific ΔΔ*G* values. The model was trained on a composite dataset that includes Megascale, FireProt, and other public sources. Since the authors did not release training code, we used the published results for comparison. Note that their training data used to train Stability Oracle partially overlaps with the test sets used in our evaluation, giving advantages to Stability Oracle compared to SPURS in the evaluation.

#### FoldX

FoldX ^4^ is a widely used physics-based method that estimates protein stability changes based on an empirical force field. Following standard protocols, we used the Repair and BuildModel steps to compute ΔΔ*G* values. In cases where the wild-type sequence of a protein did not match the amino acid sequence in the associated PDB structure due to missing residues or inconsistent annotations, FoldX raised errors for these proteins due to the unresolved inconsistency in the input structure file and mutation list. We therefore excluded these cases, which comprised a small fraction of the test variants (2,168 out of 49,790; 4%) and had only a negligible impact on the overall evaluation of FoldX’s performance.

#### ThermoNet

ThermoNet ^5^ is a structure-aware deep learning model that uses 3D convolutional neural networks on voxelized atomic neighborhoods to predict ΔΔ*G*. We used benchmark results reported by Dieckhaus et al.^1^ for benchmarking.

#### PROSTATA

PROSTATA ^6^ is a machine learning model that applies transfer learning to a pre-trained protein language model (ESM-2). It encodes wild-type and mutant sequences separately and combines their transformer-derived embeddings using regression heads to predict ΔΔ*G*. Evaluation results were obtained from Dieckhaus et al.^1^.

#### Rosetta

Rosetta ^7^ is a comprehensive physics-based modeling platform for protein structure and energetics. Stability predictions are computed using energy-based modeling of mutations. We used performance reported in Dieckhaus et al.^1^ for comparison.

#### DDGun

DDGun ^8^ is a family of untrained methods for predicting protein stability changes upon mutation. The sequence-based variant relies on multiple sequence alignments, while the structure-based version incorporates 3D structural features. We used the official implementation from https://github.com/biofold/ddgun to perform ΔΔ*G* predictions.

#### MutateEverything

MutateEverything ^9^ is a parallel decoding framework for efficient ΔΔ*G* prediction across all single and higher-order variants. The model fine-tunes an AlphaFold backbone with a lightweight MLP decoder on the Megascale dataset. We used the official implementation from https://github.com/jozhang97/Mutate Everything to perform prediction for high-order mutation evaluations.

